# Spectral organization of individualized connectome harmonics across brain structure, function and cognition

**DOI:** 10.64898/2026.07.19.739422

**Authors:** Thomas A. W. Bolton, Mikkel Schöttner Sieler, Jagruti Patel, Michael Chan, Dimitri Van De Ville, Patric Hagmann

## Abstract

Connectome harmonics provide a spectral representation of structural brain connectivity that has emerged as a powerful framework for studying structure–function relationships. However, two fundamental questions remain unresolved: how should inter-individual variability in structural connectivity be incorporated into graph signal processing analyses, and at which spectral scale should connectome harmonics be interpreted? Here, using structural and functional MRI data from 875 participants, we show that subject-specific connectome harmonics reveal a reproducible hierarchical organization of the human connectome. Across the spectrum, harmonics exhibited orderly transitions in stability, graph support, sparsity and anatomical localization, and naturally organized into multiscale families sharing common structural properties. Sparse harmonic representations accurately reconstructed brain activity across resting-state and task paradigms, revealing frequency-dependent recruitment of this structural hierarchy during cognition. Finally, combining structural and functional harmonic organization enabled the prediction of individual cognitive performance, demonstrating that both representations capture complementary behaviorally relevant information. Together, our findings show that connectome harmonics should be viewed not simply as graph-frequency modes of structural connectivity, but as a hierarchically organized representation of the connectome that links brain structure, functional dynamics and cognition.

## Introduction

Elucidating the organizational principles of the brain has remained a central objective of neuroscience ^1–5^. Structural brain architecture spans multiple spatial scales, from neurons and synapses to the cortical sheath and eventually, hundreds of structurally distinct brain regions interconnected by white-matter axonal bundles^6–13^. At the macroscale, several organizational principles supporting efficient information processing are conserved across individuals ^14–17^. This structural organization constrains the spatiotemporal patterns of brain activity^18,19^. Coordinated activity across distributed brain regions ultimately gives rise to lower- and higher-order brain functions^20–22^.

While the fundamental principles of brain organization are conserved across individuals, structural connectivity also exhibits substantial inter-individual variability (IIV)^23^. Because structural architecture constrains brain activity, this variability is expected to contribute to individual differences in functional organization and, ultimately, cognition and behavior. Indeed, IIV in structural connectivity has been associated with variability in cognitive abilities as well as with the manifestation of brain disorders ^13,24,25^. Consequently, methods that explicitly account for structural individuality should provide a richer description of brain function than approaches relying solely on population-average representations.

Magnetic resonance imaging (MRI) enables the non-invasive study of brain structure and function at the whole-brain scale. Diffusion-weighted MRI yields an estimate of whole-brain *structural connectivity* (SC), while functional MRI (fMRI) provides an indirect measure of brain activity over time. Network science^26–30^ has considerably advanced our understanding of large-scale brain organization by characterizing interactions across brain networks ^31,32^. More recently, gradient analysis ^33,34^ has provided a complementary low-dimensional description of brain organization by identifying the principal axes of structural or functional variation. Graph signal processing (GSP; see Ortega et al. ^35^, Dong et al. ^36^, Leus et al. ^37^ for reviews) naturally extends these frameworks by treating functional activity as signals evolving on a structural graph, thereby providing a unified spectral representation of brain structure and function. Within this framework, structural connectivity is decomposed into so-called *connectome harmonics*, each of which encodes signal variations across brain regions at a specific spectral graph frequency^38,39^. Harmonics with increasing graph frequency relate to more segregated and less integrated features of brain connectivity^40^, linking GSP to graph theoretical tools; at the same time, the connectome harmonics basis can itself be interpreted as a generalization of gradient representations (see Lioi et al. ^41^ for a detailed explanation). Together, harmonics define a spectral reference frame derived from structural connectivity in which functional activity can be represented at frame-wise temporal resolution, enabling the joint characterization of spectral and temporal aspects of structure–function relationships^42^.

Despite these advantages, two fundamental methodological questions remain unresolved: first, **how should inter-individual variability in structural connectivity be incorporated into the GSP framework?** Most studies derive connectome harmonics from an average or consensus structural connectome across participants, thereby providing a common spectral reference frame shared by all individuals ^43–54^. This strategy greatly facilitates comparisons across subjects, as identical harmonic indices correspond to the same spatial patterns. However, it also neglects structural variability between individuals, despite such variability being closely linked to behavioral variability ^13,24,25,55^. An alternative is to derive harmonics separately for each participant^40,56–58^, which naturally captures individual structural organization, but introduces new challenges because the resulting harmonic bases are no longer directly comparable across subjects, requiring explicit correspondence between harmonics to be established. Consequently, whether structural individuality should be treated as nuisance variability to suppress, or as meaningful information to exploit, remains an open question. Second, **at which spectral scale should connectome harmonics be interpreted?** Existing studies typically aggregate harmonics into heuristic spectral bands or select predefined frequency cutoffs^40,45,49–51,53,56,57,59,60^, yet it remains unclear whether behaviorally relevant information is organized at the level of individual harmonics, broad spectral partitions, or intermediate groups of harmonics.

Here, we address both questions using subject-specific connectome harmonics derived from individual structural connectivity. We first establish a principled framework for comparing subject-specific harmonic representations across individuals, allowing structural individuality to be incorporated into GSP analyses rather than treated as nuisance variability. Building on this correspondence, we investigate the spectral organization of behaviorally relevant information and determine whether connectome harmonics should be interpreted individually or as larger spectral entities. Using diffusion MRI from 875 participants from the Human Connectome Project, we show that although connectome harmonics are mathematically defined as individual eigenmodes, behaviorally relevant information is organized into reproducible spectral subfamilies. These families provide a more meaningful level of organization than isolated harmonics or arbitrarily selected frequency bands. Building on this structural framework, we integrate resting-state and task-based functional MRI to characterize how individualized harmonics are dynamically recruited across cognitive states and to determine whether their functional expression provides complementary information for behavioral prediction. Finally, we demonstrate that explicitly accounting for this spectral organization improves the prediction of cognitive abilities, linking structural individuality, functional dynamics and behavior within a unified GSP framework.

## Materials and Methods

### Overview of analyzed subjects and data

We analyzed data from 875 participants of the Human Connectome Project (HCP)^61^ Young Adult dataset (age: 28.67 ± 3.7 years; 412 males). For each participant, diffusion MRI, resting-state fMRI (four fifteen-minute scans), task-based fMRI (two scans for each of the seven HCP paradigms: EMOTION, GAMBLING, LANGUAGE, MOTOR, RELATIONAL, SOCIAL and WM), and behavioral assessments were available. For details regarding ethnicity or head movement, please refer to the **Supplementary Materials**.

Resting-state fMRI scans comprised four 15-minute acquisitions collected over two separate scanning days (hereafter denoted REST_1_ and REST_2_, respectively). Throughout the manuscript, REST_1_ serves as the primary resting-state dataset, whereas REST_2_ is used to assess the reproducibility of the reported findings.

For each resting-state session as well as for task-based recordings, the first scan (referred to as SES_1_) was obtained with a left-right phase encoding direction, while the second (SES_2_) instead utilized a right-left phase encoding direction. SES_1_ and SES_2_ scans were treated independently when learning and validating the proposed sparse spectral representation, thereby avoiding circularity in subsequent analyses.

To obtain a parsimonious representation of behavioral variability across participants, we used a recent decomposition of the behavioral scores provided by the HCP into four factors through exploratory factor analysis (see Schöttner et al. ^62^ for details), yielding four factors reflective of (1) *Mental health*, (2) *Cognition*, (3) *Processing speed*, and (4) *Substance use*. Of note, a highly similar decomposition was also obtained from the HCP data in a separate publication using independent component analysis ^63^, providing independent support for the robustness of the extracted behavioral dimensions.

Details regarding data preprocessing to generate structural connectivity matrices and regional activity time courses can be found in the **Supplementary Materials**.

### Generation of connectome harmonics

For each subject *s* = 1, 2*,…, S*, structural connectivity was represented by a weighted adjacency matrix **A**^(^*^s^*^)^ ∈ R*^R^*^×^*^R^*, where *R* = 216 denotes the number of cortical regions and 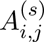 corresponds to the normalized fiber density between regions *i* and *j*. Equivalently, **A**^(^*^s^*^)^ defines an undirected weighted graph G^(^*^s^*^)^ = (*V, ε*), with |*V*| = *R* nodes and |*ε*| ≤ *R*(*R* − 1)/2 edges.

Rather than deriving connectome harmonics from a structural connectome averaged across participants, harmonics were computed independently for every subject in order to preserve inter-individual structural variability. For each subject, the combinatorial graph Laplacian was computed as 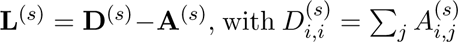. The symmetric normalized Laplacian was then obtained as 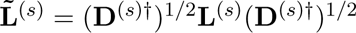, where † denotes the Moore-Penrose pseudoinverse, allowing the normalization to remain well defined in the presence of isolated nodes.

The connectome harmonics were obtained by solving 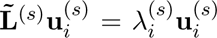, which yields **L̃**^(^*^s^*^)^ = **U**^(^*^s^*^)^**Λ**^(^*^s^*^)^**U**^(^*^s^*^)⊤^. In these equations, **U**^(^*^s^*^)^ ∈ R*^R^*^×^*^R^* is the basis of connectome harmonics for subject *s*, arranged as columns; we refer to each individual harmonic as 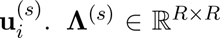 is a diagonal matrix containing the eigenvalues 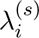 associated with each harmonic. Eigenvalues were ordered in ascending order, corresponding to increasing graph frequency, such that the first non-trivial harmonics capture smooth signal variations over the structural graph, whereas higher-order harmonics represent progressively more spatially oscillatory patterns.

Because connectome harmonics are computed independently for every participant, their ordering is not guaranteed to correspond across subjects despite being ordered according to graph frequency. The following subsection therefore introduces a framework for establishing harmonic correspondence between participants.

### Matching of harmonics across subjects

Because harmonics are computed independently for each participant, corresponding harmonics must be identified before cross-subject analyses can be performed. Following previous studies employing subject-specific connectome harmonics ^40,56,57^, harmonics were aligned across participants using the Hungarian algorithm ^64^. Pairwise assignment costs were defined as 1 − |corr(·, ·)|, with corr the spatial Pearson correlation coefficient between two harmonics from separate subjects. Absolute correlations were used because eigenvectors are defined only up to sign. Consequently, two harmonics differing only by a global sign inversion should be regarded as equivalent.

Subject 1 was chosen as the reference participant, and harmonics from all remaining participants were reordered with respect to this reference. Following harmonic assignment, matched harmonics were subsequently multiplied by ±1 whenever necessary to maximize their spatial Pearson correlation with the corresponding reference harmonic. We did not observe qualitative differences in our results when repeating the alignment using alternative reference subjects. Furthermore, the evolution of cross-subject similarity between matched harmonics across spectral frequency in our hands (see **Supplementary Figure 7A**) closely matched previous observations^40^. Examples of aligned connectome harmonics for a subset of participants are shown in **Figure 1A**.

**Figure 1:**
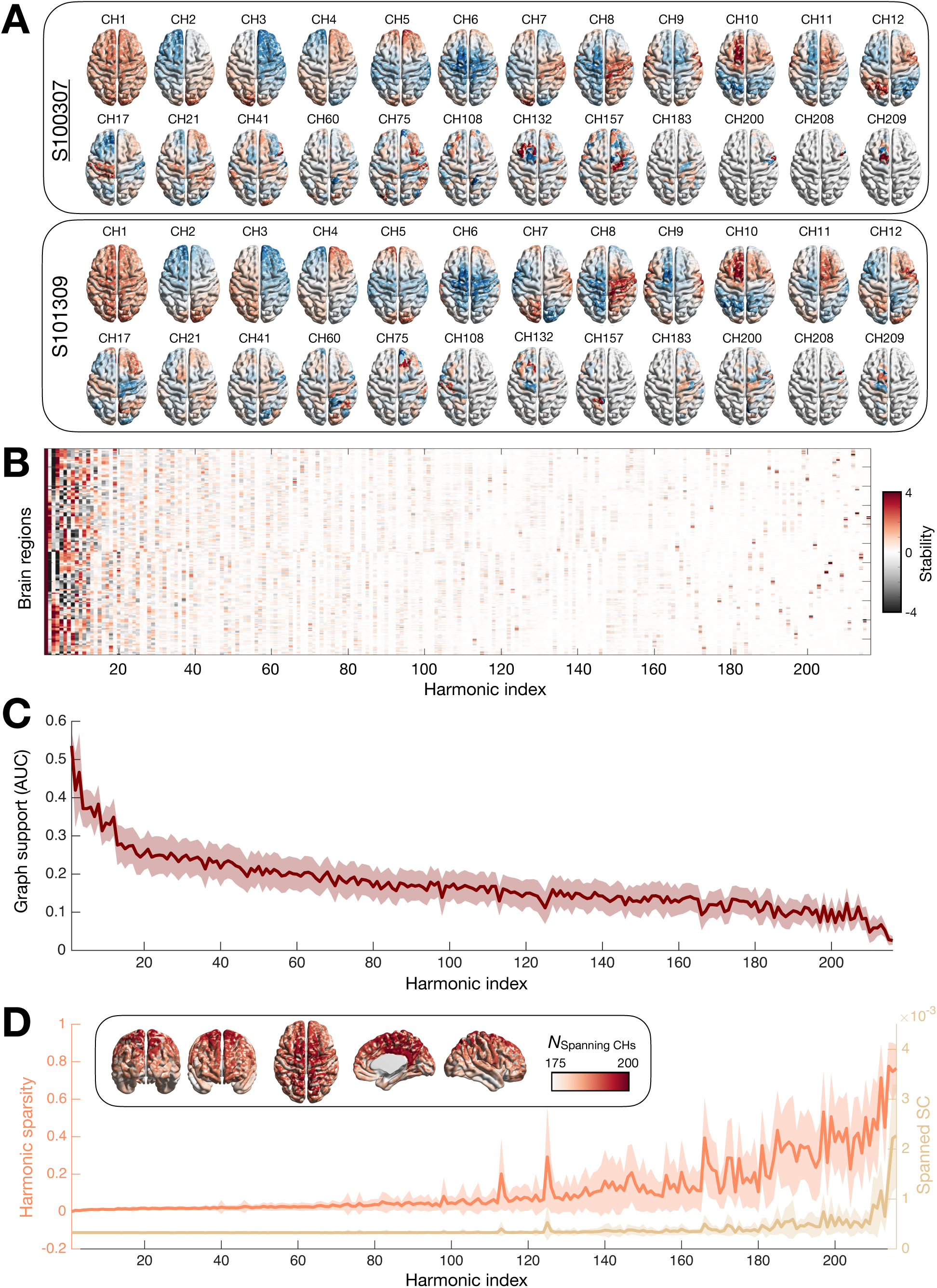
Connectome harmonics undergo progressive structural specialization across the graph frequency spectrum. **(A)** Selected matched connectome harmonics spanning the graph frequency spectrum for two representative subjects. Harmonics from the second subject were reordered to maximize correspondence with those of the first subject. **(B)** Population-wise regional stability of connectome harmonic signal, quantified as the mean divided by the standard deviation across subjects for each brain region (rows) and harmonic (columns). **(C)** Normalized graph support as a function of harmonic index. Graph support was quantified as the normalized area under the decay profile of normalized harmonic magnitude as a function of normalized graph distance from the harmonic peak. **(D)** Harmonic sparsity (orange; fraction of supra-threshold brain regions) and spanned structural connectivity (light brown; mean structural connectivity among supra-threshold regions) as a function of harmonic index. (*Inset*) Spectral participation of brain regions, defined as the number of harmonics for which a region exhibited supra-threshold signal. Throughout the figure, shaded regions denote standard deviation across subjects. Together, these analyses reveal a progressive transition from globally distributed to increasingly localized structural representations across the harmonic spectrum. CH: connectome harmonic, AUC: area under the curve, SC: structural connectivity.

### Characterization of subject-specific connectome harmonics Harmonic descriptors

To characterize how reproducible individual harmonics were across participants, we quantified the consistency of the signal expressed by each brain region within each harmonic. We devised a measure of *stability* for the signal in each region of each harmonic, quantified as the population-wise mean divided by the population-wise standard deviation. Stability increases whenever a brain region exhibits consistently large positive or negative signal values across participants while displaying limited inter-subject variability. Stability is displayed in **Figure 1B** while mean and standard deviation effects are individually shown in **Supplementary Figure 2B–C**.

To quantify normalized graph support, for each connectome harmonic, the cortical region exhibiting the largest absolute harmonic magnitude was identified. Graph distances from this peak region to every other cortical region were then computed as shortest-path distances on the individual structural connectivity graph. Harmonic magnitude was normalized by its maximum value, and graph distance was normalized by the largest observed graph distance for that harmonic. Graph support was finally defined as the normalized area under the resulting decay profile of harmonic magnitude as a function of graph distance, computed using the trapezoidal rule. Larger graph support therefore indicates harmonic patterns spanning broader graph neighborhoods, whereas smaller values correspond to harmonics confined to progressively smaller structural subnetworks.

Following Sipes et al., harmonic coefficients whose absolute value was smaller than 0.001 were considered negligible ^40^. We then computed *sparsity* as the fraction of regions with non-null signal, and *spanned structural connectivity* (sSC) as the average structural connectivity between all retained region pairs. Let all regions surviving thresholding for harmonic *k* in subject *s* be part of set *P_k_*, then 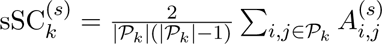. Additionally, we quantified how many harmonics spanned each brain region. Further complementary harmonic descriptors, as well as the impacts of edge weight and graph topology on sparsity, are addressed in the **Supplementary Materials**.

### Structural connectivity reconstruction

To visualize the structural connectivity represented by individual harmonics, we reconstructed approximate structural connectomes following Sipes et al. ^40^. A harmonic 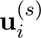 and its associated eigenvalue 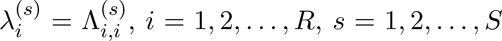, were used to generate 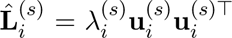, the rank-one approximation of the normalized Laplacian using only harmonic *i*. The underlying approximate SC was then obtained as 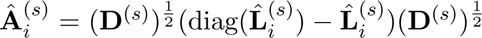, setting negative-valued elements to zero and normalizing with respect to the Frobenius norm. In the **Supplementary Materials**, we examine the evolution of subsequently derived graph-theoretical measures across spectral frequency. Note that the formulation naturally extends to any subset of harmonics, by summing the corresponding rank-one contributions before reconstructing the structural connectivity matrix.

### Cross-subject similarity assessments

To gain insight into how similar matched harmonics were across subjects, we first considered the evolution of similarity between matched harmonics (Pearson’s correlation coefficient) as a function of spectral frequency (**Supplementary Figure 7A**). We then generalized this to all possible harmonic pairs: similarity between harmonic *i* from subject *k* and harmonic *j* from subject *l* was defined as 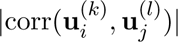. The absolute value is required because only matched harmonics undergo explicit sign alignment.

To identify groups of harmonics sharing similar cross-subject similarity profiles, the columns of the harmonic similarity matrix were first z-scored and subsequently submitted to agglomerative hierarchical clustering (weighted average linkage method, cosine distance). The number of clusters was selected from the dendrogram together with the block structure of the similarity matrix, yielding *F* = 19 spectral families. To quantify the validity of the resulting family organization, we compared the mean pairwise similarity between harmonics belonging to the same family with the mean pairwise similarity between harmonics belonging to different families. Statistical significance was assessed by permutation testing, randomly shuffling family labels while preserving family sizes. Family assignments were subsequently used only for visualization and interpretation, whereas all prediction analyses relied directly on the continuous similarity values.

Family-wise properties were studied through the same predictors as in harmonic-wise analyses. Additionally, we also quantified family size, spectral span (distance in harmonic indices between the first and last included harmonics), and spectral centroid (center of gravity in harmonic index). **Supplementary Table 1** summarizes these results.

### Structural prediction framework

To evaluate whether subject-specific connectome harmonics capture behaviorally relevant structural information, we predicted the *Mental health*, *Processing speed*, *Cognition* and *Substance use* factors reported by Schöttner et al. ^62^. Predictions were performed using kernel ridge regression (KRR), following previous benchmarking studies of behavioral prediction from connectivity data ^65^. Predictions were based on structural connectomes reconstructed from progressively larger subsets of low-frequency harmonics. Specifically, for each value of *k_L_* ∈ [2*, R*], we reconstructed an approximate structural connectivity matrix using the first *k_L_* connectome harmonics, following the procedure described above. This enabled us to quantify how prediction performance evolved as increasingly broader portions of the harmonic spectrum were incorporated into the structural representation. For the *Cognition* factor, an analogous analysis was performed by progressively reconstructing structural connectivity from the highest-frequency harmonics, allowing the predictive contribution of both ends of the spectrum to be evaluated.

To characterize behavioral prediction at the level of harmonic families, we used a similar approach to predict *Cognition* (1) by gradually accumulating families from the one with the smallest mean spectral index, and (2) by predicting from individual entities only. Prediction improvements upon the addition of a new family 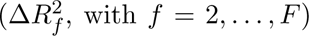, and prediction performance using individual families 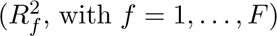, were examined in conjunction to delineate subtypes of families.

Prediction performance was quantified using the coefficient of determination *R*^2^ within a nested cross-validation framework comprising 100 random outer 80/20 train-test splits, and an inner 3-fold cross-validation procedure for hyperparameter optimization. Cross-validation folds respected the HCP family structure, such that related participants were never split across training and test sets. The regularization parameter was optimized over 400 logarithmically spaced candidate values between 10^−4^ and 10^4^. To account for potential confounding effects of head motion, predictions were repeated both with and without mean framewise displacement included as a covariate of no interest.

### Sparse spectral representation of functional activity

Let **X**^(^*^s^*^)^ ∈ R*^R^*^×^*^T^* denote the regional fMRI activity recorded for subject *s*, where *R* = 216 is the number of brain regions and *T* the number of acquired fMRI volumes. The activity at time point *t* is denoted by 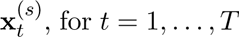. In the classical graph Fourier transform (GFT)^38,39^, functional activity is represented in the connectome harmonic basis as **X̂**^(*s*)^ = **U**^(^*^s^*^)⊤^**X**^(^*^s^*^)^. Rather than using this dense representation, we sought sparse representations of brain activity in the connectome harmonic basis.

Sparse representations were obtained using the SPAMS toolbox ^66^. For each participant, the subject-specific connectome harmonics **U**^(^*^s^*^)^ were used directly as the dictionary atoms, such that the dictionary remained fixed throughout the analyses and only the sparse coefficients were estimated.

Let 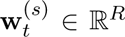 denote the sparse coefficient vector describing activity at time *t*. Coefficients were estimated by solving:

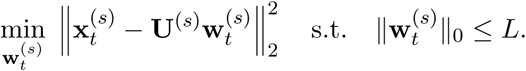

Sparse coefficients were estimated independently for every fMRI time point. In practice, this formulation enforces that only a small subset of connectome harmonics contributes to the reconstruction of each instantaneous brain activity pattern, thereby yielding a temporally resolved sparse spectral representation of brain activity. The collection of all coefficient vectors then forms 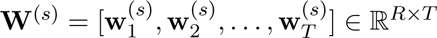. Each coefficient *W_r,t_* quantifies the instantaneous expression strength of harmonic *r* at time *t*. Because connectome harmonics are ordered by increasing graph frequency, low-index coefficients describe smooth structural modes, whereas high-index coefficients capture progressively more segregated modes.

The same sparse coding procedure was applied independently to resting-state and task-based fMRI recordings, yielding one sparse spectral representation for every acquisition and participant. The optimal sparsity level L was selected exclusively from the SES_1_ recordings. Candidate values ranging from 1 to *R* were evaluated, and reconstruction RMSE was computed for each sparsity level. The optimal sparsity level, denoted *L*^∗^, was defined as the knee point of the resulting root mean square error curve, corresponding to the best compromise between reconstruction accuracy and representation sparsity. This fixed sparsity level was subsequently used to estimate sparse representations for the independent SES_2_ recordings, ensuring that all downstream analyses were performed on data independent from the sparsity selection procedure.

### Dynamic metrics of harmonic expression

Sparse coding naturally partitions the temporal evolution of each harmonic into periods of activity and inactivity, according to whether its corresponding sparse coefficient is non-zero. Note that we consider a harmonic *active* when it is either positively or negatively expressed. In the **Supplementary Materials**, we conduct validation analyses that separately examine positive and negative activations.

Let *τ_k_* denote the set of time points when each harmonic *k* was active; we then quantified the *intensity* of activation as 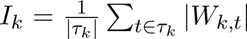. Additionally, we computed two metrics reflective of harmonic temporal dynamics inspired by dynamic functional connectivity analyses^67^. *Entry rate* was defined as the number of transitions from the inactive to the active state per unit time, whereas *duration* corresponded to the average length of contiguous active periods.

Intensity, entry rate, and duration provide complementary descriptions of harmonic temporal dynamics. To obtain a composite metric and streamline some of our analyses, we explicitly quantified signal power. To reduce the influence of head motion, harmonic power was computed only over active time points whose framewise displacement remained below 0.5 mm ^68^, denoted by *τ̃_k_*. We then computed harmonic power as 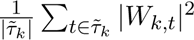.

All metrics were obtained for the SES_2_ recording of each paradigm, for each connectome harmonic of each participant. Averaging these metrics across participants enabled us to characterize the spatiotemporal organization of harmonic expression across the spectral axis and identify distinct regimes of harmonic dynamics. Comparisons across paradigms were subsequently used to investigate how harmonic expression relates to cognitive function.

Because entry rate and duration would be strongly altered by the exclusion of scrubbed time points, entry rate, duration and intensity were quantified on non-scrubbed recordings. In the **Supplementary Materials**, we explicitly verified that associations with head movement remained minimal.

### Statistical comparison of harmonic dynamics across tasks

To explore whether intensity, entry rate and duration were behaviorally relevant, we contrasted each measure individually across the different paradigms from the HCP dataset. More specifically, we focused on the SES_2_ scans from the seven task-based HCP paradigms and the REST_1_ acquisition, while the REST_2_ SES_2_ data was utilized as a negative control, by explicitly verifying (1) the absence of REST1 vs REST2 differences, and (2) the presence of similar REST vs task patterns across both REST conditions.

First, separately for each metric and harmonic, we performed a one-way repeated-measures ANOVA with task as factor (8 levels, one per paradigm). To assess the robustness of the results to potential motion-related confounds, these analyses were repeated in an extended model including mean framewise displacement, together with the task × mean framewise displacement interaction term.

Next, to further characterize significant task effects, we performed post-hoc ANCOVAs for all 28 task contrast pairs, separately for each metric and harmonic. Each model included task (2 levels), mean framewise displacement (continuous covariate), and their interaction. As an additional robustness analysis, we also evaluated all pairwise contrasts using the non-parametric Wilcoxon signed rank test, without explicit modeling of head movement.

All resulting *p*-values were Bonferroni-corrected across the complete set of parallel tests (3 metrics × 28 task contrasts × 216 harmonics). Significant results were subsequently examined to determine which dynamic metrics differed between task conditions and how these differences were distributed across the connectome harmonic spectrum.

### Hybrid structure–function prediction framework

To evaluate whether functional harmonic dynamics provide complementary information to structural harmonics for behavioral prediction, we constructed three prediction models: (i) a structure-only model based exclusively on connectome harmonic similarity, (ii) a function-only model based exclusively on harmonic expression dynamics, and (iii) a hybrid model jointly combining structural and functional similarity. Predictions were performed using kernel ridge regression within the same nested cross-validation framework described above.

Harmonic power was used as the functional descriptor. Only the seven task paradigms together with the REST_1_ acquisition were included (*N_T_* = 8), to avoid over-representing resting-state recordings. 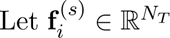 the vector of power values across *N_T_* = 8 paradigms for harmonic *i* and subject *s*. The similarity between subjects *k* and *l* is defined as:

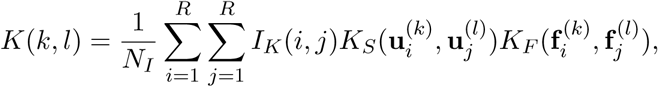

where *NI* = Σ*_i,j_ IK*(*i, j*) normalizes the kernel by the number of included harmonic pairs.

The kernel evaluates pairs of harmonics across subjects according to their structural similarity, 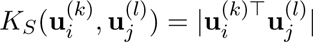, and their functional similarity, 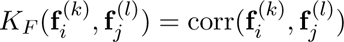.

Setting the indicator matrix *I_K_* to the identity restricts the comparison to matched harmonics only, thereby assuming perfect harmonic correspondence across subjects. Conversely, setting it to a matrix of ones allows all cross-harmonic comparisons and explicitly incorporates the family organization of connectome harmonics. Finally, fixing either *K_F_* or *K_S_* to a constant yields the structure-only and function-only kernels, respectively.

We compared structure-only, function-only and hybrid models, each with and without accounting for similarity between non-matched harmonic pairs. Statistical significance of prediction performance was assessed by comparing the observed *R*^2^ values against null distributions generated from 100 random permutations of the behavioral scores.

To benchmark the proposed harmonic framework against conventional connectivity representations, prediction performance was also evaluated using whole-brain structural connectivity and functional connectivity matrices. FC features consisted of vectorized functional connectivity matrices concatenated across paradigms, whereas SC features corresponded to vectorized structural connectivity matrices. Subject similarity was quantified using Pearson correlation, yielding baseline SC and FC kernels for comparison with the harmonic-based models.

In all prediction analyses, mean framewise displacement computed over the retained (non-scrubbed) frames was included as a covariate of no interest, since harmonic power was estimated exclusively from these frames. Supplementary analyses additionally evaluated prediction performance without including this confound.

### Statistical analyses

All statistical analyses were performed in MATLAB R2025b. Statistical procedures are described within their corresponding methodological subsections. Unless otherwise specified, statistical significance was defined as *p <* 0.05 after correction for multiple comparisons whenever applicable.

## Results

### Frequency-dependent structural specialization of connectome harmonics

Representative harmonics revealed a gradual transition from smooth, spatially distributed patterns at low graph frequencies to increasingly localized and heterogeneous patterns at higher frequencies (**Figure 1**). Low-frequency harmonics showed strongly conserved regional expression patterns across individuals, although subject-specific differences in amplitude and spatial distribution remained visible, whereas intermediate-frequency harmonics exhibited substantially greater inter-individual variability despite the harmonic matching procedure. Greater similarity was observed once more for some of the highest-frequency harmonics. This observation suggests that although intermediate-frequency harmonics capture the greatest structural individuality, highest-frequency harmonics recover reproducible fine-scale structural organization. Signal sparsity also increased progressively with graph frequency, such that connectome harmonics gradually transitioned from smooth, spatially distributed patterns to increasingly localized representations.

We quantified these observations through a measure of population-wise stability in regional harmonic signal (**Figure 1B**). Regional stability remained uniformly high throughout the lowest-frequency portion of the spectrum before progressively declining with graph frequency. Intermediate-frequency harmonics showed a mix of almost-null and low stability values across regions, reflecting increased sparsity and inter-subject variability in harmonic signal. High-frequency harmonics became even sparser, but also showed small subsets of stable regions. Non-parametric statistical testing confirmed a gradually increased sparsity, but also showed that significantly stable regions remained in the intermediate frequency band (**Supplementary Figure 2D**).

Next, we asked which properties from the structural connectivity matrices were associated with this organization. We quantified the normalized graph support of each connectome harmonic as the normalized area under the decay profile of harmonic magnitude with increasing graph distance from its peak magnitude (**Figure 1C**). Graph support gradually decreased along the frequency spectrum (Spearman’s correlation coefficient with harmonic index: *R* = −0.98,*p <* 10^−6^), indicating that increasing graph frequency is associated with progressively smaller graph neighborhoods, providing a structural basis for the emergence of sparse harmonic patterns.

Furthermore, the progression of sparsity along the frequency spectrum (**Figure 1D**) closely tracked that of localization (Pearson’s correlation coefficient across subjects: *R* = 0.97 ± 0.02), indicating that sparse harmonic representations remain confined to local neighborhoods in the Euclidean distance sense; and of spanned structural connectivity (*R* = 0.77 ± 0.06), suggesting that structural connectivity strength contributes to the emergence of sparsity. Notably, sparsity started increasing before spanned structural connectivity, suggesting that sparse harmonic expression is not solely a consequence of connectivity weights, but also reflects changes in graph organization. Follow-up analyses (**Supplementary Figure 4**) confirmed this observation, showing that sparsity of the harmonics is abolished both when weights are not considered anymore, and when graph topology is lost.

We also examined how frequently each region exhibited suprathreshold harmonic signal across harmonics (**Figure 1D**, inset). Spectral participation was highest along the cortical midline, progressively decreasing when moving towards the frontal, occipital or temporal lobes. Regions participating in a larger number of connectome harmonics preferentially occupied connector roles within the structural connectome (**Supplementary Figure 6**). Spectral participation was most strongly associated with betweenness centrality (*R* = 0.26 ± 0.08) and participation coefficient (*R* = 0.21 ± 0.09), whereas only modest associations were observed with strength (*R* = 0.14 ± 0.12) and virtually no relationship emerged with clustering coefficient (*R* = 0.02 ± 0.1) or eigenvector centrality (*R* = 0.05 ± 0.1). These findings suggest that harmonic diversity is preferentially supported by connector regions mediating communication between structural modules, rather than by highly connected or locally clustered regions.

### Connectome harmonics organize into reproducible spectral families carrying behavioral information

Given that individual harmonics progressively diverged in their structural organization across the spectrum, we next asked whether this divergence followed a reproducible higher-order organization.

Cross-subject similarity between matched harmonics (**Supplementary Figure 7A**) followed a U-shape profile, remaining high for the first 5% of the spectrum, then gradually decreasing. It remained low throughout the intermediate-frequency range before increasing again among the highest-frequency harmonics. This highlights that the most conserved harmonics across subjects lie at both extremes of the spectrum, fully in line with previously reported observations^40^.

Extending the analysis to all pairwise cross-subject similarities (**Figure 2A**) revealed a clear block structure along the diagonal at low graph frequencies, indicating groups of neighboring harmonics sharing similar cross-subject patterns (as annotated by vertical orange lines). With increasing frequency, these blocks progressively broadened while their internal similarity decreased, before eventually dissolving into sparse isolated similarity peaks at the highest graph frequencies. Although the block structure suggested an organization into families of harmonics, it remained unclear whether similarity was purely driven by spectral proximity. We therefore performed hierarchical clustering directly on harmonic similarity profiles, revealing a multi-scale organization of connectome harmonics (**Supplementary Figure 8**). For subsequent analyses, we retained the partition into 19 families, corresponding to the first resolution at which the prominent low-frequency similarity blocks became individually resolved.

**Figure 2:**
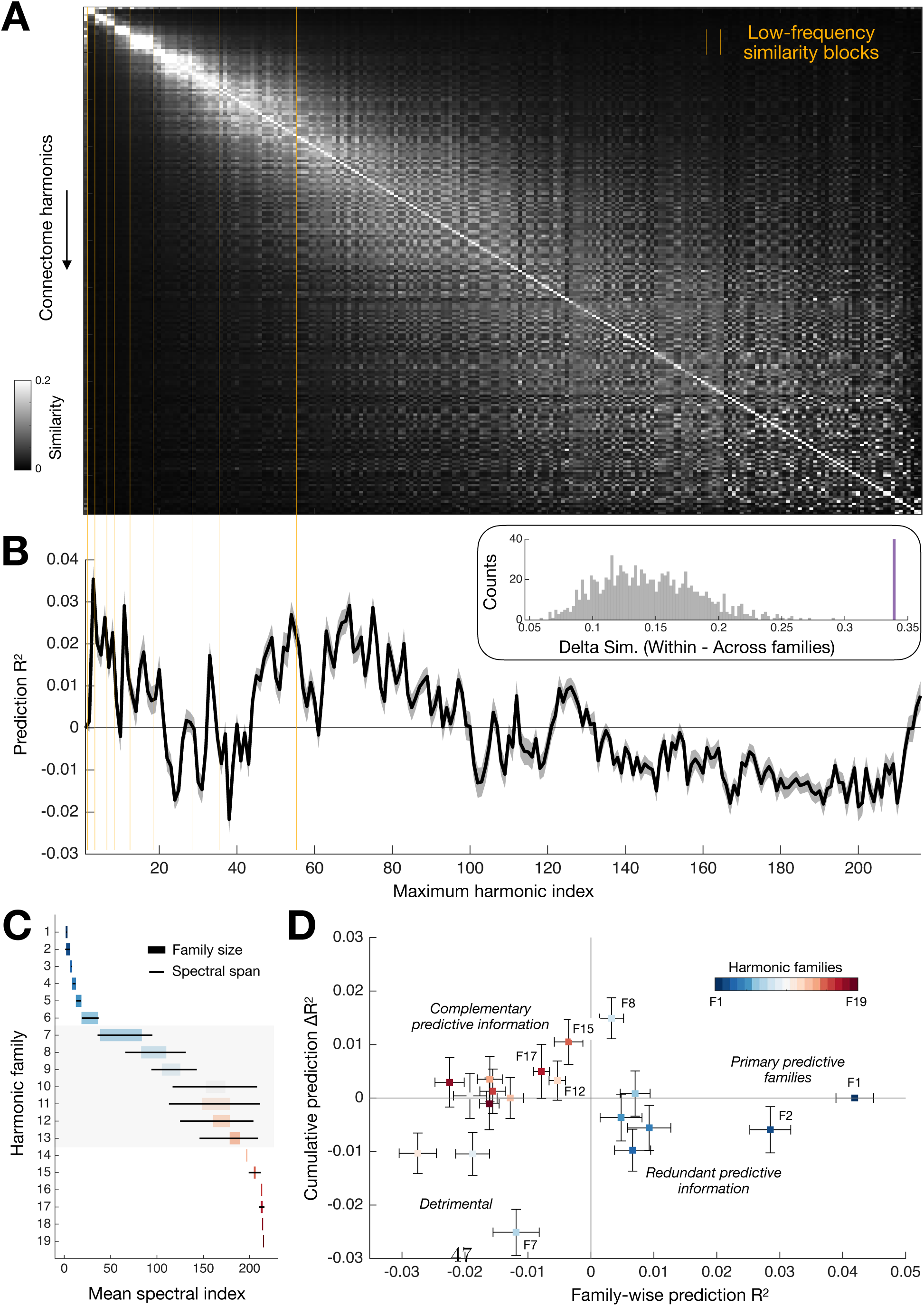
Connectome harmonics organize into reproducible spectral families that differentially contribute to cognition. **(A)** Average cross-subject similarity across all pairs of connectome harmonics, computed as the absolute dot product. Vertical orange bars delineate visually apparent low-frequency similarity blocks. The color bar is saturated at a similarity of 0.2 to provide an exhaustive view of how similarity organizes across the spectrum. (*Inset*) Difference between within-family and across-family similarity after hierarchical clustering of harmonic similarity profiles (purple bar), compared with the corresponding null distribution (grey histogram). **(B)** Coefficient of determination *R*^2^ obtained on left-out data, in a nested cross-validation scheme, when predicting the *Cognition* factor score using structural connectivity approximations reconstructed with cumulatively increasing numbers of harmonics (X-axis), starting from the lowest-frequency ones. Shaded regions denote standard error of the mean. **(C)** For all 19 extracted harmonic families, rectangle width denotes family size, while horizontal black bars indicate spectral span. Color-coding denotes family index. Families without a visible spectral span consist of a single harmonic. **(D)** Coefficient of determination 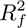 when predicting *Cognition* from structural connectivity approximated by an individual family *f* (X-axis), and the corresponding change in prediction performance 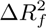 obtained when family *f* is added during cumulative family-wise prediction (Y-axis). Color-coding denotes family index, and error bars denote standard error of the mean. Subtypes of families with distinct behavioral relevance are annotated. Together, these results identify harmonic families as coherent multiscale units carrying distinct behavioral information. Sim.: similarity, F: family.

To verify whether this organization was robust, we computed average similarity within and across families, and took their difference as statistic of interest. Under a null hypothesis accounting for block sizes and spectral contiguity, within-minus cross-family similarity difference was strongly significant (Δ*_Sim_*=0.339, *p <* 10^−6^; see **Figure 2B** inset and **Supplementary Figure 9**).

In terms of spectral organization, family size remained low at both extremes of the spectrum, including only between 2 and 6 harmonics; however, it became much larger in the middle, where between 18 and 45 harmonics were included in the same family. Families identified by hierarchical clustering were not necessarily spectrally contiguous, as spectral span and spectral dispersion increased markedly for intermediate-frequency families (**Figure 2C**, **Supplementary Table 1**), demonstrating that harmonics with similar cross-subject organization may be separated along the graph frequency spectrum.

Family-wise properties followed the same sequential organization observed for individual harmonics, with stability declining before graph support, followed by rises in sparsity, and then in anatomical localization and spanned structural connectivity (**Supplementary Figure 10**). Thus, the hierarchical organization identified at the harmonic level extends naturally to harmonic families. To determine whether this spectral organization carried behavioral information, we examined how well subsets of harmonics with increasing spectral bandwidth could predict *Mental health*, *Processing speed*, *Cognition* and *Substance use*. Successful prediction could only be achieved for Cognition (**Figure 2B**, see **Supplementary Figure 11** for the other scores). Prediction accuracy exhibited three distinct spectral regimes, occurring when only the lowest-frequency harmonics were retained, when harmonics were accumulated up to the low-to-intermediate frequency regime, and, to a lesser extent, after inclusion of the full spectrum. Annotating the boundaries between visible blocks of harmonic cross-similarity revealed that changes in prediction accuracy frequently coincided with the completion of a subset, suggesting that behaviorally relevant information is organized at the level of harmonic families rather than isolated harmonics.

Including mean framewise displacement as a covariate preserved the shape of the prediction curve, yielding slightly larger prediction accuracy (**Supplementary Figure 12**). Furthermore, when adding harmonics from the highest-frequency one, the obtained curve differed markedly (**Supplementary Figure 13**): prediction became significant when reaching intermediate-frequency harmonics, but then stabilized without ever rising to the same level as for the main analysis, indicating that indiscriminate inclusion of higher-frequency harmonics does not improve prediction and may dilute behaviorally relevant information carried by lower-frequency modes.

The coincidence between changes in prediction accuracy and the completion of harmonic families motivated a complementary family-wise analysis, where harmonics were accumulated family by family according to increasing mean spectral index. Prediction accuracy evolved non-monotonically across the spectrum, with successive family additions producing alternating improvements and degradations in cognitive prediction (**Supplementary Figure 14A**). At the global level, the lowest and highest-frequency families were particularly salient, and a rise in prediction accuracy was also observed in the middle of the spectrum.

Quantifying the incremental contribution of each family by computing the change in prediction accuracy (Δ*R*^2^) following the addition of each successive family revealed that a subset of intermediate- and high-frequency families produced the largest positive Δ*R*^2^ values (**Supplementary Figure 14B**, top half). On the contrary, family 7 was particularly deleterious to prediction, whereas neighboring families exerted markedly different effects.

We complemented these observations by training prediction models on individual families, to dissociate family-specific contributions from interactions with previously incorporated families (**Figure 2D** and **Supplementary Figure 14B**, bottom half). The first two low-frequency families (F1 and F2) yielded substantial standalone prediction of cognition, acting as primary predictive families, while further accumulation of low-frequency families (F3-F6) produced no increase in performance despite small positive contributions when taken individually, reflective of redundant predictive information. Family 7 degraded performance both when considered in isolation or in a cumulative manner, an example of a family detrimental to the prediction of cognition. In contrast family 8, together with a subset of high-frequency families (F12, F15 and F17), were more beneficial to prediction when considered in the cumulative analysis than on their own, highlighting the fact that they provide complementary predictive information not captured by the dominant low-frequency representation.

These results indicate that behaviorally predictive information is unevenly distributed across harmonic families, with specific families providing disproportionate contributions. The contribution of intermediate- and high-frequency families appears to depend on the preceding spectral representation rather than on independent behavioral information, suggesting that cognition is encoded hierarchically across harmonic families.

In summary, connectome harmonics are organized into reproducible spectral families that undergo a progressive transition from globally distributed, stable representations to localized structural motifs. These families constitute reproducible multiscale building blocks of the harmonic spectrum that jointly encode behaviorally relevant structural information.

### Connectome harmonics exhibit frequency-dependent temporal dynamics organized into two dynamical regimes

Having established that connectome harmonics undergo frequency-dependent structural specialization and organize into reproducible families, we next investigated whether these structural differences were reflected in their temporal expression during ongoing brain activity. We derived sparse spectral representations of fMRI signals for every recording across all HCP paradigms. **Figure 4A** (top panel) showcases regional time courses of activity for an indicative resting-state recording.

**Figure 3:**
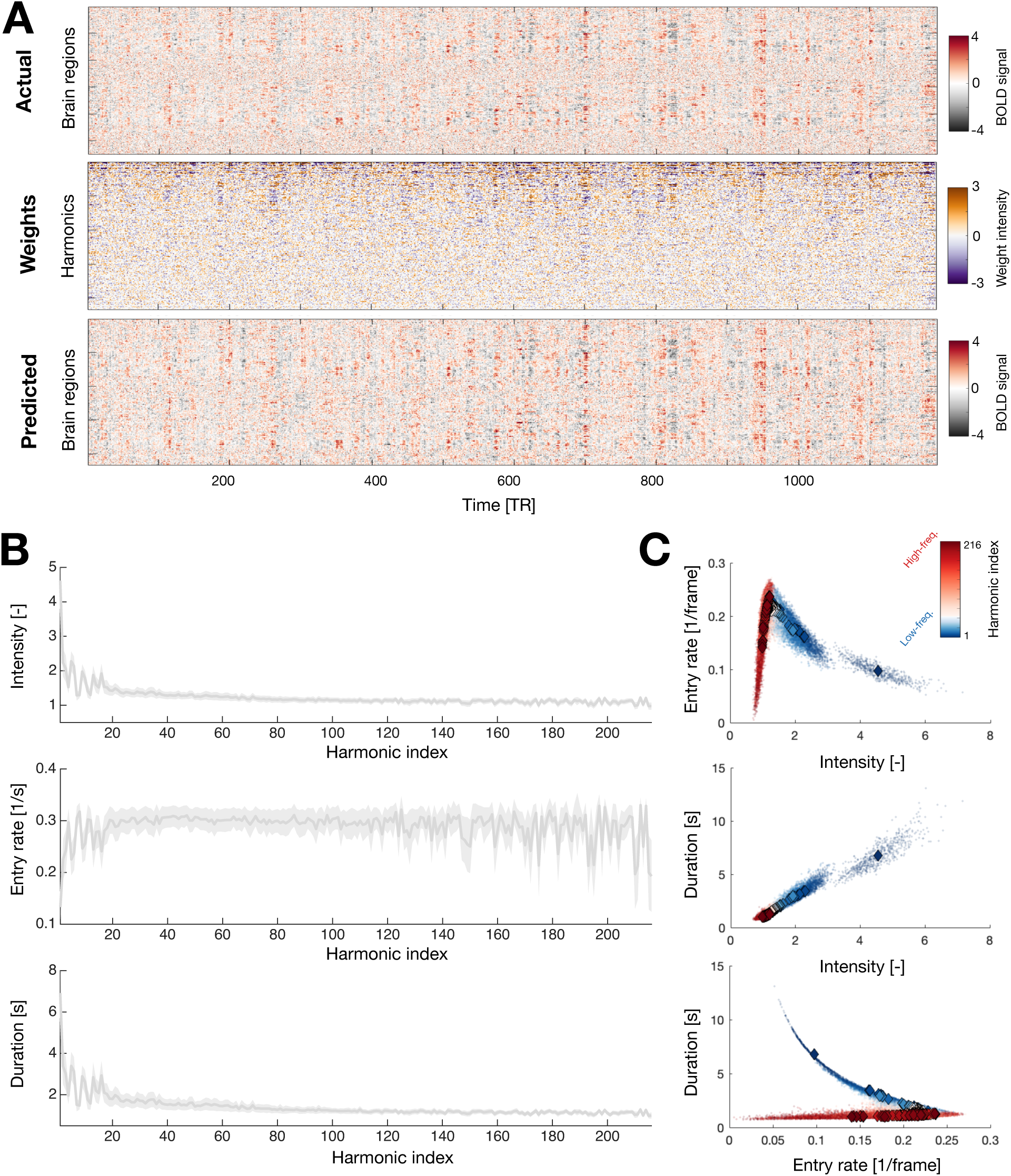
Connectome harmonics exhibit sparse temporal dynamics organized into two frequency-dependent dynamical regimes. **(A)** (*Top*) Indicative regional time courses of activity for a selected REST2 recording. (*Middle*) Corresponding connectome harmonic expression weights, obtained upon sparse spectral decomposition with cross-validated regularization parameter. (*Bottom*) Reconstructed regional activity time courses obtained from the sparse harmonic representation. **(B)** Variation across the harmonic spectrum of the intensity (top), entry rate (middle) and average duration (bottom) of harmonic expression. Shaded regions denote standard deviation across subjects. **(C)** JRelationships between pairs of temporal metrics across the harmonic spectrum (color-coded by harmonic index). Diamonds indicate population-average values for each harmonic, whereas dots represent individual subjects. Together, these analyses reveal a continuous transition between two frequency-dependent dynamical regimes, from low-frequency harmonics exhibiting strong, sustained expression to high-frequency harmonics characterized by weak, transient burst-like activations.

**Figure 4:**
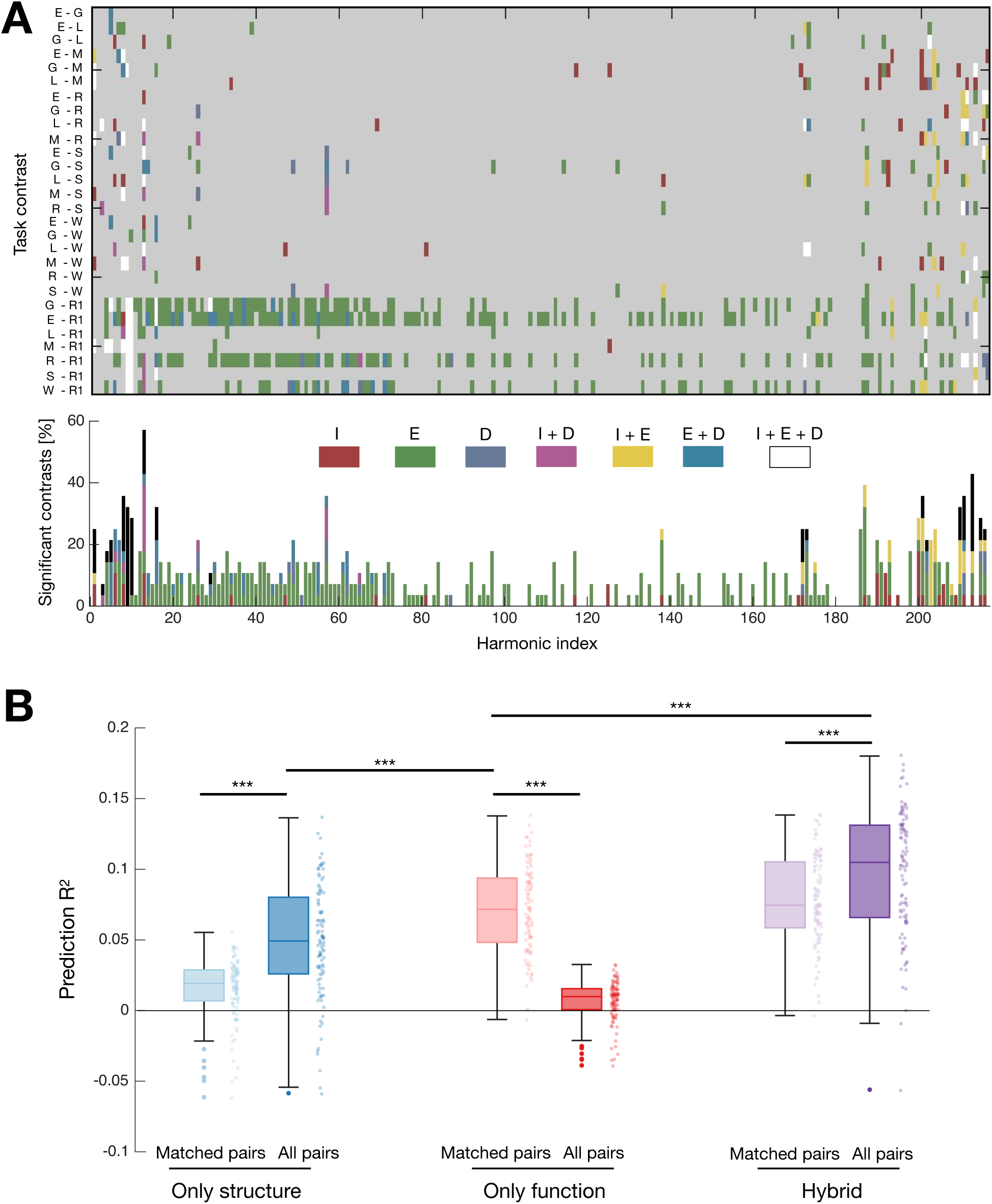
Task-dependent harmonic dynamics complement structural harmonic organization in predicting cognition. **(A)** (*Top*) For each task contrast (rows) and harmonic (columns), the color indicates which combination of temporal metrics reached significance. Grey entries denote non-significant comparisons after Bonferroni correction. (*Bottom*) Percentage of significant task contrasts for each harmonic, with color-coding the same as above. **(B)** Coefficient of determination *R*^2^ for the prediction of *Cognition* using kernels constructed from structural similarity between harmonics (blue), functional similarity between cross-task harmonic power profiles (red), or their combination (purple). Light box plots correspond to kernels computed from matched harmonic pairs only, whereas dark box plots include all harmonic pairs. Mean framewise displacement computed on fMRI volumes surviving scrubbing at 0.5 mm ^68^ was included as a covariate of no interest. Together, these analyses demonstrate that structural relationships across the harmonic spectrum and task-dependent harmonic dynamics provide complementary information for predicting cognition. ***: *p <* 0.001. I: intensity, E: entry rate, D: duration.

The middle panel shows the associated expression weights for all harmonics, where low-frequency harmonics (top rows) exhibit the largest fluctuations in expression weight. The bottom panel displays predicted time courses of activity. Prediction quality was excellent across the board, with median *R*^2^ values reaching 0.89 for all considered paradigms (**Supplementary Figure 16**). Despite this excellent reconstruction accuracy, only 39–41% of the harmonic spectrum was expressed at any given time point on average (**Supplementary Figure 17**), demonstrating that ongoing brain activity occupies only a restricted portion of the available harmonic space.

We next quantified three complementary properties of harmonic expression: average intensity, entry rate (the fraction of time points at which a harmonic became active), and average duration of expression. **Figure 4B** displays these quantities across graph frequencies for the example case of REST_2_ recordings, while **Supplementary Figures 18–20** show equivalent results for all other paradigms. Global properties of the examined features were similar across cases: expression intensity and duration were maximal for the lowest-frequency harmonics, but did not decrease monotonically with spectral frequency, as evidenced by local irregularities in the curves. This non-monotonic behavior likely reflects the distinct large-scale graph partitions represented by successive low-order eigenmodes rather than simple ordering by graph frequency. A progressive decrease was then seen as graph frequency increased. Entry rate increased rapidly among the lowest-frequency harmonics before reaching a plateau of approximately 0.3*s*^−1^, with greater fluctuations reappearing at the highest graph frequencies. These behaviors were also observed when analyzing only positive or negative expression weights (**Supplementary Figure 21**), and all three metrics exhibited only weak associations with head movement (|*R*| *<* 0.2 across tasks, harmonics and metrics; **Supplementary Figure 22**).

Further examination of the relationships between these three quantities (**Figure 3C**) revealed that harmonic dynamics occupy a continuous trajectory linking two distinct dynamical regimes. Low-frequency harmonics (blue) were characterized by strong, sustained activations occurring infrequently, whereas high-frequency harmonics (red) were expressed through weaker but substantially more transient events. Intermediate-frequency harmonics formed a smooth continuum between both extremes, indicating that harmonic dynamics evolve progressively across the spectrum rather than forming discrete temporal classes. Unlike the structural similarity profiles, temporal expression metrics did not exhibit clear family-wise clustering, but instead evolved smoothly along graph frequency. Together, these observations reveal a frequency-dependent organization of harmonic dynamics that parallels the progressive structural specialization described in **Figures 1–2**.

### Task-dependent harmonic dynamics reveal complementary structural and functional information

Having characterized the intrinsic structural and temporal organization of connectome harmonics, we next asked whether these temporal expression properties are modulated by cognitive state. Specifically, we tested whether harmonic intensity, expression duration and entry rate differed systematically across resting-state and the seven HCP task paradigms. This analysis aimed to identify frequency bands whose temporal dynamics are preferentially engaged by distinct cognitive demands.

To quantify paradigm-dependent modulation while accounting for head movement, we performed one-way ANCOVAs with task as factor and mean framewise displacement together with its interaction with task as covariates, independently for each harmonic and each temporal metric. Significant task effects were distributed across the spectrum (**Supplementary Figure 23A**), but their prevalence differed markedly across metrics. Entry rate was affected most strongly (181 significant harmonics after Bonferroni correction), whereas duration and intensity exhibited considerably fewer significant task effects (71 and 91 harmonics, respectively). Motion effects were primarily seen for intensity (**Supplementary Figure 23B**; 149 harmonics against 33 and 30 for entry rate and duration, respectively). In contrast, significant task-by-motion interactions (**Supplementary Figure 23C**) were rare (11, 11 and 6 harmonics for intensity, entry rate and duration, respectively), indicating that motion influenced harmonic dynamics similarly across paradigms. We also verified that the qualitative profile of task effects, characterized by a subset of well located peaks, was insensitive to the inclusion of head movement as a covariate (**Supplementary Figure 23D**). We followed up with assessing all individual task contrasts. **Supplementary Figure 24** displays the resulting signed F-statistics, obtained by multiplying each F-statistic by the sign of the corresponding contrast estimate, while **Figure 4A** offers a summarizing view where color-coding denotes which subsets of metrics were significant across task comparisons and harmonics. In line with the above results, entry rate emerged as the dominant source of task-related modulation. More specifically, entry rates were frequently higher during task conditions than during resting-state. Overall, task-related differences were not uniformly distributed across the harmonic spectrum, but concentrated near the two extremities of the spectrum, mirroring the high cross-subject similarity and behavioral relevance previously observed for these harmonics.

An interesting parallel can also be made with the two previously identified dynamical regimes: indeed, the low-frequency side of the spectrum tended to highlight opposite signs between estimated task effects for entry rate and duration/intensity, while on the high-frequency end, all signs often coincided (**Supplementary Figure 24**).

To ascertain the specificity of our approach, we examined how similar task-versus-rest findings were between the REST_1_ and REST_2_ recordings (**Supplementary Figure 25**). No significant differences were detected between both resting-state scans for either metric and harmonic, and patterns of F-statistics to HCP tasks were strongly similar. Furthermore, similar spatial patterns were observed when analyses were repeated without motion correction and using non-parametric rank-sum testing, although substantially more harmonic-task combinations reached significance (**Supplementary Figure 26**).

Together, these observations indicate that behavioral state primarily regulates when connectome harmonics are recruited rather than how strongly or how long they are expressed.

Having established that connectome harmonic dynamics are selectively modulated across behavioral paradigms, we next asked whether *structural harmonic organization* and *functional harmonic dynamics* carry complementary information. To this end, we compared prediction accuracy on the *Cognition* score obtained from kernels derived exclusively from structural similarity, exclusively from functional similarity, or from their combination (**Figure 4B**). We considered harmonic power as a composite functional feature accounting for the joint impacts of intensity, entry rate and duration. Cross-subject similarity kernels were then constructed by aggregating structural similarity (computed as in **Figure 2**), functional similarity (defined as the correlation between harmonic power profiles across tasks), or the product of both (hybrid approach), either using only matched harmonic pairs or all possible harmonic pairings.

Using only structural similarity, significant prediction was achieved (permutation *p*-values*<* 0.01; see **Supplementary Figure 27** for corresponding null distribution), indicating that the geometry of the harmonic space itself contains behavioral information. When using only functional similarity, significant prediction was achieved as well (permutation *p*-values*<* 0.01 for matched-pairs, *<* 0.02 for all-pairs), highlighting the fact that meaningful information is also embedded in the temporal expression profiles of harmonics. Most importantly, combining structural and functional similarity yielded the highest prediction accuracy (permutation *p*-values*<* 0.01), demonstrating that both representations contribute complementary rather than redundant information.

Prediction improved when extending the structure-only analysis to all pairs of harmonics (*p <* 0.001), consistent with the presence of meaningful information beyond matched pairs only as revealed in **Figure 2**. However, prediction in the function-only case rather worsened upon broadening the set of harmonic pairings (*p <* 0.001), suggesting that while structural information generalizes across neighboring harmonics, functional information is much more harmonic-specific. Optimal performance was therefore achieved when the family-wise structural organization of connectome harmonics was incorporated into the comparison of their temporal dynamics, corresponding to the hybrid all-pairs kernel.

We verified that the results were insensitive to the inclusion of head movement as a covariate of no interest (**Supplementary Figure 28**). Furthermore, using raw structural connectivity to build the kernel produced, as expected, a prediction accuracy comparable to that of the structure-only all-pairs approach (**Supplementary Figure 29A**). Using functional connectivity concatenated across all tasks produced the highest prediction accuracy (**Supplementary Figure 29A**), fitting with its status as prediction gold standard ^65^. Interestingly, although functional connectivity yielded the highest prediction accuracy, the FC-derived kernel was largely orthogonal to the hybrid harmonic kernel (**Supplementary Figure 29B**), indicating that both approaches capture distinct sources of information.

Together, these findings indicate that connectome harmonics provide a common representation through which structural organization and functional dynamics become directly comparable. Structural similarity captures relationships that generalize across harmonic families, whereas functional similarity reflects harmonic-specific temporal recruitment. Their combination therefore provides complementary information that cannot be recovered from either modality alone.

## Discussion

Traditionally, connectome harmonics have been interpreted as graph-frequency modes of structural connectivity, whose principal characteristic is the frequency with which they oscillate across the graph ^42,43,60^. Our results suggest that connectome harmonics are more completely characterized by the joint consideration of *graph frequency* and *graph support*. While graph frequency determines how rapidly a harmonic varies across the connectome, graph support determines the spatial extent over which this variation is expressed.

This perspective naturally explains the progressive emergence of sparsity, anatomical localization, and increasing modularity/decreasing efficiency across the spectrum. Rather than representing independent phenomena, these properties can be interpreted as arising from the gradual contraction of harmonic support toward increasingly localized graph neighborhoods. Connectome harmonics can therefore be understood as a hierarchy of structural motifs spanning progressively smaller portions of the connectome.

The spatial organization of connectome harmonics has received comparatively less attention^40^, as previous work primarily emphasized their functional interpretation^43,60^, behavioral relevance ^45,49^ or reliance on long-range connectivity^52,69,70^. Our findings suggest that graph support provides a unifying spatial descriptor that links several structural properties previously considered separately. More generally, signal analysis often benefits from complementary descriptions of frequency content and spatial (or temporal) localization. Likewise, our findings suggest that connectome harmonics are most naturally interpreted through the complementary descriptors of graph frequency and graph support. Interestingly, the progressive contraction of graph support suggests that connectome harmonics§ provide an intrinsic multiscale decomposition of structural connectivity. This raises the possibility that, in the connectomic setting, some analyses traditionally approached through graph wavelet constructions^42,71^ may also be addressed by exploiting the natural hierarchy of harmonic support. From this perspective, connectome harmonics occupy an intermediate position between classical graph Fourier modes and graph wavelets, simultaneously providing a spectral ordering and a progressively localized multiscale representation.

Interestingly, this reinterpretation also offers a different perspective on inter-individual variability. Although intermediate-frequency harmonics exhibited the greatest variability across subjects, normalized graph support remained remarkably stable across the graph frequency spectrum. This suggests that graph support represents the spatial scale of structural motifs (conserved across individuals), whereas inter-individual variability may primarily reflect differences in the structural realization of motifs sharing a common spatial scale. This realization may involve both differences in local connectivity weights and shifts in the anatomical location of harmonic support.

Viewed through the joint lens of graph frequency and graph support, connectome harmonics provide a multiscale decomposition of structural connectivity that bridges global representations of brain organization with graph-theoretical analyses of the complete connectome. More broadly, this view places connectome harmonics within a growing family of low-dimensional coordinate systems for describing large-scale brain organization, alongside cortical gradients^16,33^, geometric eigenmodes ^52,69^, and low-dimensional dynamical representations^72^.

Connectome harmonics constitute an orthogonal mathematical basis of structural connectivity. Accordingly, individual harmonics are typically regarded as the elementary units of graph spectral analysis. However, our results demonstrate that harmonics are not organized as isolated biological entities, but instead cluster into reproducible families with shared structural and behavioral properties. In other words, biological organization emerges above the level of individual harmonics. Previous studies have largely treated individual harmonics as the elementary units of analysis ^43,44,46,48,54^ or, conversely, aggregated broad spectral bands into coarse frequency classes^40,45,49,53,60^. Our results reveal an intermediate, data-driven level of biological organization that naturally partitions the harmonic spectrum into structurally and behaviorally coherent families. The diverse contributions of individual families to behavioral prediction, ranging from complementary to redundant or even detrimental, support the view that they constitute functionally distinct spectral entities rather than arbitrary spectral groupings.

Beyond providing a parsimonious representation of the spectrum, harmonic families reveal that graph frequency alone does not fully capture the internal organization of connectome harmonics. The observations that individual families may encompass spectrally dispersed harmonics and exhibit a nested multiscale organization suggest that graph frequency alone is insufficient to describe the internal architecture of the harmonic spectrum.

Our results further suggest that biologically meaningful information may be distributed across neighboring harmonics within a family rather than being confined to a single eigenmode. Indeed, we have shown that even upon harmonic matching, biologically meaningful signal is distributed across harmonics belonging to the same family. This observation may also inform approaches such as connectome gradient analysis, where individual gradients (equivalent to individual harmonics^41^) are typically analyzed separately. Our results raise the possibility that biologically meaningful variation may instead extend across families of neighboring harmonics.

Finally, harmonic families offer a principled, data-driven way of defining graph spectral kernels for graph signal processing ^35,38,71^. Rather than selecting arbitrary spectral boundaries, future graph signal processing approaches may exploit the intrinsic organization of the harmonic spectrum revealed by family structure.

Structural connectivity matrices are highly similar across individuals (*R >* 0.9 in the majority of cases), yet we have seen that their harmonic decompositions exhibit substantial variability. This indicates that edge-wise similarity and similarity of the harmonic representation are fundamentally different notions.

The harmonic representation exhibits a structured pattern of variability across the spectrum. Low- and high-frequency harmonics showed comparatively greater reproducibility across subjects, whereas intermediate frequencies exhibited the largest variability. Importantly, behavioral prediction indicates that individual differences remain informative throughout the spectrum, suggesting that reproducibility and individuality coexist across graph frequencies.

One interpretation consistent with our findings is that the biologically reproducible entities may not be individual harmonics, but rather the spectral subspaces approximated by harmonic families. From this perspective, the harmonic families identified here reflect structural motifs that are conserved across subjects, whereas the exact distribution of structural information among individual harmonics within a family constitutes a subject-specific realization of that common organization. Harmonic families may therefore represent the organizational form through which inter-individual variability is expressed.

Under this interpretation, harmonic matching should not be viewed as recovering exact one-to-one harmonic correspondences, but rather as providing an approximate alignment between subject-specific realizations of a shared structural organization. Accordingly, harmonic families should be viewed not merely as empirical clusters, but as observable approximations of the conserved spectral organization shared across individuals.

This departs from most neuroimaging studies, where variability is generally addressed in predefined measurements. Subject-specific connectome harmonics elevate inter-individual variability from differences within a common representation to differences in the representation itself. Population-shared organization persists at the family level, while information can redistribute among individual harmonics across subjects. Reproducibility and variability therefore coexist at different levels.

The particularly large families observed at intermediate graph frequencies may arise from the combination of reduced eigengaps previously reported in this spectral range^40^, which increase the sensitivity of individual eigenvectors to structural perturbations, together with the richer mesoscale organization of structural connectivity^73^, which may naturally require representation across multiple neighboring harmonics.

This interpretation suggests that subject-specific harmonic organization should be regarded as a biologically meaningful object of study rather than merely a source of variability. It is also consistent with the behavioral analyses presented here. The superior performance of all-pairs prediction, together with the marked changes observed upon completion of harmonic families, suggests that behaviorally relevant structural information is distributed across harmonic families rather than confined to individual harmonics.

Future work could explicitly characterize these shared spectral subspaces and their subject-specific realizations, potentially providing a principled framework for studying individual differences in brain organization.

The Laplacian eigendecomposition provides the brain with a multiscale repertoire of structural modes characterized by graph frequency, graph support and family organization. However, the eigendecomposition itself does not prescribe how these modes should evolve over time. Our results show that the brain recruits this structural repertoire according to a highly organized temporal hierarchy, in which harmonic expression intensity, duration and entry rate vary systematically across the graph spectrum.

Rather than revealing two discrete classes of dynamics, the harmonic spectrum exhibits a continuous transition linking persistent recruitment of broadly supported structural modes to weaker, short-lived recruitment of increasingly localized modes. This organization provides a common framework within which two perspectives that have often been treated separately in systems neuro-science—continuous spatiotemporal activity^72,74,75^ and transient event-based descriptions of brain dynamics ^76–79^—can coexist.

Interestingly, the harmonic spectrum is highly asymmetric: a relatively small number of low-frequency modes account for much of the overall expression intensity, whereas a substantially larger repertoire of higher-frequency harmonics contributes through individually weaker, transient recruitment. Brain activity therefore appears to rely simultaneously on a compact set of strongly expressed global modes that account for much of the overall signal, and a substantially richer repertoire of localized transient modes that expand its dynamical flexibility.

Interestingly, this temporal organization is not limited to a simple transition from sustained to transient recruitment. The joint evolution of expression intensity, duration and entry rate suggests qualitatively different modes of recruitment across the spectrum. Low-frequency harmonics combine strong expression with long durations and infrequent entries, consistent with recurrent, sustained engagement of broad structural modes. By contrast, the highest-frequency harmonics exhibit both short durations and low entry rates, indicating that they are recruited as comparatively rare and spatially localized events rather than through continuous rapid switching. Intermediate graph frequencies form a continuum linking these two extremes.

The observed temporal hierarchy suggests a coupling between structural scale and temporal persistence. Broadly supported harmonics tend to be recruited strongly and persistently, whereas increasingly localized harmonics are recruited through weaker, shorter-lived events. Because our analyses do not model the temporal evolution of the Laplacian itself, this coupling should not be interpreted as a mathematical consequence of graph frequency or graph support. Rather, it represents an empirical property of how the brain recruits the structural harmonic basis.

One possible mechanistic explanation for this temporal hierarchy is that broad structural modes facilitate the integration of activity across distributed cortical systems and therefore favor more sustained recruitment, whereas localized modes can support more spatially confined computations that are assembled and dissolved over shorter timescales. Under this view, the observed temporal hierarchy would reflect a coupling between the spatial scale of structural organization and the characteristic timescale over which that organization is recruited. Whether this coupling reflects the propagation of activity across large-scale structural pathways, recurrent network interactions, or other mechanisms remains to be established.

Although only approximately 40% of harmonics are expressed at any individual time point, every harmonic contributed to the sparse representation at some point during the recording. The harmonic spectrum therefore appears not to consist of “important” and “irrelevant” modes. Instead, different harmonics participate on different temporal scales, collectively enabling a sparse instantaneous representation while fully exploiting the structural repertoire over time. The observation that only a subset of harmonics is expressed at any individual time point, despite the eventual recruitment of the full harmonic repertoire, also relates to the broader question of the effective dimensionality of large-scale brain activity^80–83^. Rather than suggesting that only a restricted portion of the harmonic basis is biologically relevant, our results indicate that the brain dynamically explores different portions of a much richer structural repertoire over time.

The overall temporal organization remained remarkably similar across all behavioral paradigms, indicating that the multiscale structural hierarchy imposes a canonical mode of harmonic recruitment. Cognitive demands did not abolish this organization, but selectively redistributed harmonic expression in a reproducible, harmonic-dependent manner. In other words, the structural hierarchy defines a baseline dynamical scaffold, upon which cognition induces systematic yet comparatively modest reallocations of harmonic recruitment.

An intriguing possibility is that temporal recruitment may itself be organized at the level of harmonic families rather than individual harmonics, with the harmonic-wise dynamics reported here representing subject-specific realizations of a conserved higher-order organization that may involve coordinated patterns of recruitment extending beyond individual harmonics. Resolving this question will require methodological developments capable of directly characterizing the dynamics of spectral subspaces.

Together, Figures 2 and 3 suggest that both the structural organization of the harmonic spectrum and its temporal recruitment are intrinsic properties of the connectome harmonic framework. Cognitive processes therefore appear to act primarily by selectively modulating pre-existing multiscale structural and dynamical organizations, rather than by creating entirely new modes of organization.

Having established that connectome harmonics constitute a multiscale structural and dynamical organization, the remaining question is whether this organization carries behaviorally meaningful information. Our results indicate that cognition is best captured by jointly considering the structural organization of subject-specific harmonics and their functional recruitment across behavioral paradigms. Structural similarity alone and functional similarity alone each contributed complementary information, while their combination consistently yielded the best prediction performance. Rather than treating structure and function as independent modalities, these findings suggest that cognition is more faithfully represented by how function recruits an individualized structural harmonic scaffold.

Our findings also have methodological implications for graph signal processing. Previous work has largely focused on identifying informative functional descriptors or appropriate graph spectral filters. Our results suggest that improving the structural representation is equally important. Accounting for the organization of harmonics into families through all-pairs structural similarity systematically improved prediction, indicating that biologically meaningful information extends beyond one-to-one harmonic correspondences. At the same time, the comparatively weaker performance of functional all-pairs similarity suggests that structural organization provides a biologically meaningful constraint that limits the otherwise large solution space of possible functional similarities. Future developments may therefore benefit from jointly refining both the structural representation (for example through family- or support-based kernels) and the functional representation (by combining multiple temporal descriptors or exploiting richer measures of harmonic similarity).

Although functional connectivity achieved higher prediction performance than the present hybrid representation, this does not diminish the relevance of the proposed framework. Functional connectivity directly summarizes statistical interactions between brain regions and therefore represents a natural upper benchmark for many prediction tasks. By contrast, the objective of the present work was to determine whether connectome harmonics provide an informative structural-functional coordinate system in which cognition can be represented. The complementary contributions of structural and functional harmonic information, together with the consistent above-chance prediction achieved by the hybrid representation, support this interpretation.

Taken together, our results suggest that cognition is not encoded by isolated harmonics or individual temporal metrics, but rather emerges from the coordinated recruitment of a multiscale structural organization. Structure defines the harmonic repertoire, dynamics govern how this repertoire is recruited, and cognition selectively modulates this pre-existing organization. This perspective naturally unifies the principal findings of the present work and provides a foundation for future graph signal processing approaches that explicitly integrate subject-specific structural organization with functional dynamics.

### Limitations

First, our analyses were restricted to the cerebral cortex. Although cortical networks capture much of the large-scale organization of human brain function, extending the present framework to subcortical and cerebellar structures^63,84^ will be important to determine whether the multiscale organization described here generalizes to a more complete representation of brain connectivity, and whether it broadens the range of accessible brain-behavior associations.

Second, our conclusions were derived from connectome harmonics computed using the normalized graph Laplacian. Whether graph support, harmonic families and temporal recruitment emerge similarly when alternative graph operators^85^ or spectral embeddings^86^ are employed remains to be established.

Third, although harmonic families consistently emerged across the population and provide a coherent interpretation of inter-individual variability, they currently represent empirically identified clusters rather than explicitly identified invariant spectral subspaces. Developing methods capable of explicitly identifying conserved spectral subspaces represents an important direction for future work.

Fourth, the temporal hierarchy described here should not be interpreted as directly reflecting neuronal timescales or specific physiological mechanisms. The observed coupling between structural scale and temporal recruitment was established at the level of BOLD dynamics and will require validation using complementary acquisition modalities and computational models. In addition, we focused on MRI-derived measures of brain structure and function. Whether the multiscale organization identified here extends to higher-temporal-resolution modalities such as MEG or EEG (modalities in which graph signal processing has also recently gained traction ^46,47,50^) remains an important question, particularly given the temporal hierarchy reported in Figure 3.

Fifth, the present characterization of harmonic recruitment relies on a sparse representation of regional activity. Although this framework accurately reconstructed brain activity and yielded reproducible temporal organization, alternative representations of harmonic dynamics may reveal complementary aspects of functional recruitment.

### Conclusion

Together, our findings suggest that connectome harmonics should no longer be viewed simply as graph-frequency modes of structural connectivity. Instead, they constitute a multiscale, hierarchically organized representation of structural connectivity. Graph support complements graph frequency at the level of individual harmonics, harmonic families reveal an additional level of spectral organization, and task-dependent expression dynamics demonstrate how this structural organization is recruited to support cognition. Connectome harmonics should therefore be viewed not simply as graph Fourier modes, but as the building blocks of a multiscale structural, dynamical and cognitive organization of the human connectome.

## Acknowledgements

Data were provided by the Human Connectome Project, WU-Minn Consortium (Principal Investigators: David Van Essen and Kamil Ugurbil; 1U54MH091657) funded by the 16 NIH Institutes and Centers that support the NIH Blueprint for Neuroscience Research; and by the McDonnell Center for Systems Neuroscience at Washington University. This work has been supported by Swiss National Science Foundation grant #197787.

The authors acknowledge the use of ChatGPT (OpenAI GPT-5.5) during the preparation of this manuscript. The model was used as an interactive scientific writing assistant to discuss the interpretation of results, refine the conceptual organization of the manuscript, improve the clarity and structure of the Discussion, and provide stylistic feedback on the text. All scientific analyses, methodological developments, implementation, interpretation of the final results, and editorial decisions were performed and validated by the authors, who take full responsibility for the content of the manuscript.

We are grateful to Oscar Esteban, Yasser Alemán Gómez and Silas Forrer for insightful discussions regarding several parts of this work.

## Data availability

The neuroimaging data analyzed in this study were obtained from the Human Connectome Project (HCP) S1200 release and are publicly available to qualified researchers through the Human Connectome Project under the HCP Open Access Data Use Terms. Derived data supporting the findings of this study are available from the corresponding author upon reasonable request. Source data underlying the figures are provided with this paper.

## Code availability

The MATLAB code used to compute individualized connectome harmonics, perform the analyses, and generate the figures presented in this study will be made publicly available via a GitHub repository and archived on Zenodo upon publication.

## Author contributions

T.B. and P.H. designed the study. M.S. processed the resting-state and task-based functional MRI data. J.P. processed the diffusion-weighted MRI data. M.C. and D.V.D.V. provided guidance throughout the development of the work. T.B. performed all analyses and wrote the manuscript, which was reread by all authors.

