## Supplementary Information for "Spectral organization of individualized connectome harmonics across brain structure, function and cognition"

---

This Supplementary Information contains **Supplementary Methods**, **Supplementary Results**, **Supplementary References**, 4 **Supplementary Tables**, and 28 **Supplementary Figures**.

---

45 pages · 4 **Supplementary Tables** · 28 **Supplementary Figures**

**Supplementary References** **13**

**Supplementary Tables** **14**

**Supplementary Figures** **18**

### Supplementary Methods

*Magnetic resonance imaging data preprocessing*

All MRI data were obtained from the minimally preprocessed HCP release and comprised T1-weighted, diffusion-weighted, resting-state functional and task-based functional MRI scans. The additional preprocessing steps performed in this study for the generation of structural connectomes and regional functional activity time series are detailed below.

**Generation of structural connectomes.** *Connectome Mapper 3* (CMP3) RC4[1,2] was used to compute the structural connectomes, using minimally preprocessed T1-weighted and diffusion MRI data as input. For segmentation of the T1-weighted images, *FreeSurfer*[3] recon-all version 6.0.1 was used, and parcellation was done with the Lausanne 2018 atlas[4–8] at scale 3, for a total of $R = 216$ cortical brain regions entering the analyses.

Whole-brain tractograms were reconstructed with *MRtrix*[9] using deterministic tractography from white-matter seeds, constrained spherical deconvolution (maximum spherical harmonic order = 8), and 10 million output streamlines. The tractograms and parcellated anatomical images were subsequently combined within CMP3 to generate weighted structural connectomes, where edge weights corresponded to normalized fiber density.

**Extraction of regional activity time courses.** Minimally preprocessed functional MRI data had already undergone distortion correction, motion correction, registration to structural images, bias-field correction, spatial normalization and brain masking as part of the HCP preprocessing pipeline. Additional preprocessing was performed using *FSL*[10] together with in-house Python scripts (Python 3.10.9). For each scan, the first six functional volumes were discarded to prevent scanner drift. The Lausanne 2018 atlas was subsequently warped from diffusion space to Montreal Neurological Institute (MNI) space using *FSL*'s `applywarp` command with nearest-neighbour interpolation. Confound regression included the six rigid-body motion parameters together with their temporal derivatives, as well as the mean white-matter and cerebrospinal-fluid signals, all obtained from the HCP release. Voxelwise detrending, temporal high-pass filtering (0.01 Hz), and confound regression were performed simultaneously using `nilearn.clean_img`, thereby ensuring that nuisance regression remained orthogonal to temporal filtering and avoiding the reintroduction of artifacts[11]. Volumes with FD exceeding 0.5 mm were flagged for motion scrubbing and excluded from all subsequent analyses relying on scrubbed functional data[12]. Finally, regional time series were extracted using `NiftiLabelsMasker.fit_transform` (*nilearn*), after which each regional time series was standardized by temporal z-scoring.

#### *Harmonic alignment*

To realign harmonics across subjects, the MATLAB function `munkres` (Munkres' Assignment Algorithm, Modified for Rectangular Matrices, http://csclab.murraystate.edu/bob.pilgrim/445/munkres.html) was used on MATLAB R2025b. To verify that harmonic alignment was not driven by the choice of reference subject, key analyses were repeated using several arbitrarily selected reference subjects. The qualitative conclusions remained unchanged across all tested reference choices.

#### *Sparse coding implementation*

The sparse representation of harmonic expression over time was estimated with MATLAB's `mexOMP` routine from the *SPAMS* toolbox[13] through orthogonal matching pursuit. The algorithm was constrained to select at most $L$ connectome harmonics per time point. Because connectome harmonics form an orthonormal basis, all dictionary atoms had unit $\ell_2$-norm and no additional normalization was required.

For each participant and each paradigm independently, root mean square error was evaluated as $\mathrm{RMSE}_L^{(s)} = \sqrt{\frac{\left\|\mathbf{X}^{(s)} - \mathbf{D}^{(s)}\mathbf{W}_L^{(s)}\right\|_F^2}{RT}}$, where $\mathbf{X}^{(s)}$ is the activity data for subject $s$ for a $\mathrm{SES}_1$ scan, $\mathbf{D}^{(s)}$ is the dictionary made of the connectome harmonics from the same subject, and $\mathbf{W}_L^{(s)}$ contains the sparse expression coefficients obtained when constraining the representation to at most $L$ active harmonics per time point, with $L = 1, \ldots, R$. RMSE values across all candidate sparsity levels were then assembled into a participant-specific RMSE curve.

The knee point of the participant-specific RMSE curve was identified using MATLAB's `knee_pt` function (D. Kaplan, Knee Point, MATLAB Central File Exchange, retrieved July 2, 2026). The resulting sparsity level $L^*$ was subsequently used for sparse coding of the independent $SES_2$ recordings.

#### *Dynamic metric computation*

Entry rate and average duration of harmonic expression were computed using the `CAP_ComputeMetrics` function from the MATLAB *TbCAPs* toolbox[14]. Harmonic activity was binarized by considering all non-zero sparse coefficients as active time points.

To separately characterize positive and negative harmonic expression, the sparse coefficient matrices were partitioned according to coefficient sign. For analyses restricted to positive expression, negative coefficients were set to zero. Conversely, for analyses restricted to negative expression, positive coefficients were discarded and the sign of the remaining coefficients was inverted, such that expression magnitude remained directly comparable across the two cases. Dynamic metrics were then computed independently using the same procedure as in the main analyses.

#### *Hybrid kernel implementation*

For each prediction model, the complete subject-by-subject kernel matrix was computed directly from the formulation described in the main text. Kernel entries were evaluated independently for every pair of participants by averaging the selected structure–function similarity products over the harmonic pairs specified by the indicator matrix $\mathbf{I_K}$. No additional kernel normalization or scaling was applied beyond the normalization by $N_I$, such that kernel values directly reflected the average similarity across the retained harmonic pairs.

Because both structural similarity $K_S$ and functional similarity $K_F$ are symmetric, all kernel matrices were symmetric by construction. Depending on the considered model, $\mathbf{I_K}$ was either set to the identity matrix (matched harmonics only) or to a matrix of ones (all harmonic pairs). Structure-only and function-only kernels were obtained by fixing $K_F$ or $K_S$ to a constant value of one, respectively.

### Supplementary Results

The following supplementary analyses provide additional validation of the principal findings reported in the main manuscript and further characterize the structural, spatial, dynamical and behavioral organization of individualized connectome harmonics.

*Extended demographics*

**Supplementary Table 1** provides an extended overview of the demographic characteristics of the study cohort, including participant age, sex, ethnicity, and race.

*Figure 1. Structural properties of individualized connectome harmonics*

To further characterize the structural organization encoded by subject-specific connectome harmonics, we quantified their spectral, spatial and graph-theoretical properties, and evaluated their robustness to alternative graph constructions.

**Spectral organization. Supplementary Figure 1** illustrates the evolution of graph eigenvalues across the harmonic spectrum. Eigenvalues increase smoothly with harmonic index, indicating that weighted structural connectivity gives rise to a continuous spectrum of graph frequencies rather than exhibiting naturally occurring spectral boundaries.

**Origin of graph support. Supplementary Figure 2** further characterizes the stability measure introduced in the main manuscript by decomposing it into the mean (panel B) and standard deviation (panel C) of harmonic coefficients across subjects, and by identifying statistically significant stability values (panel D). Average coefficients remained close to zero across most of the spectrum, indicating that graph support is not driven by systematic shifts in harmonic amplitudes. Instead, increasing graph support reflects the progressive concentration of harmonic coefficients around zero for an increasing number of brain regions, while a progressively smaller subset of regions retains consistently non-zero values. Statistical testing confirmed that this spatial organization becomes increasingly pronounced toward higher graph frequencies.

**Spatial organization. Supplementary Figure 3** depicts the evolution of net zero-crossing (following Sipes et al.[15]), sparsity (discussed in the main material), and localization, defined as the inverse of the average pairwise distance between retained regions after thresholding. The evolutions of net zero-crossing and sparsity along the frequency spectrum are in line with previous results on independent datasets[15]. Localization increases markedly during the final quarter of the spectrum, closely paralleling the evolution of spanned SC presented in **Figure 1C**. Together, these observations indicate that high-frequency harmonics contrast regions that remain strongly structurally connected while becoming increasingly confined to localized portions of the connectome.

**Robustness to graph construction.** To determine whether harmonic sparsity depends primarily on connection weights, network topology, or both, connectome harmonics were recomputed after selectively disrupting each property (**Supplementary Figure 4**).

When only connection weights were removed by binarization (blue curves), the resulting harmonics exhibited a steeper eigenvalue spectrum but completely abolished harmonic sparsity, demonstrating that edge weights are required for sparse harmonic representations. Degree-preserving randomization (green curves) likewise eliminated sparsity and yielded nearly uniform eigenvalues, indicating that meaningful graph topology is also necessary for structured harmonic organization. Combining both perturbations (purple curves) produced similar behavior, with reduced variability across eigenvalues because the absence of edge weights further constrains the spectrum.

Together, these results indicate that sparse harmonic organization emerges from the joint contribution of biologically meaningful edge weights and network topology, rather than from either property alone.

**Graph-theoretical characterization of reconstructed structural connectomes.** To further characterize the structural organization encoded by individual harmonics, graph-theoretical measures were computed from the reconstructed harmonic-wise structural connectivity matrices[16] (**Supplementary Figure 5**). Modularity increased almost linearly with harmonic index (linear regression model, $R^2 = 0.86$, model: $Q = 0.64782 + 0.26312x$), while efficiency showed an initial plateau followed by a more rapid decay ($R^2 = 0.9$, model: $E = -1.278 \cdot 10^{-7}x^2 + 9.31 \cdot 10^{-6}x + 0.005233$). Higher-frequency harmonics reflected progressively more modular and less efficient structural organization, consistent with previous observations[15]. These observations extend previous work[15] by showing that the transition from integrated to segregated structural organization is continuous across the harmonic spectrum, revealing a gradual reorganization of structural topology across graph frequencies rather than abrupt transitions between predefined spectral regimes.

**Graph-theoretical characterization of graph support.** To better understand the structural determinants of graph support, we examined its relationships with several classical graph-theoretical measures computed from individual structural connectivity matrices (**Supplementary Figure 6**). Spectral participation was positively associated with all investigated metrics, but showed its strongest relationships with betweenness centrality and participation coefficient, indicating that graph support primarily reflects the extent to which brain regions participate in the integration of information across distributed structural networks. These findings provide additional biological intuition for the concept of graph support introduced in the main manuscript.

*Figure 2. Harmonic families and behavioral prediction*

**Reproduction of previously reported cross-subject similarity patterns.** Before investigating harmonic families, we first examined whether the cross-subject similarity structure reported in previous work could be reproduced in the present dataset (**Supplementary Figure 7**). Despite substantial differences in cohort size, preprocessing and methodological choices, both studies exhibited the same characteristic U-shaped evolution of cross-subject harmonic similarity across the graph spectrum. The pairwise similarity matrices likewise revealed comparable large-scale organization, providing independent confirmation that the similarity structure underlying harmonic families represents a robust property of individualized connectome harmonics rather than a dataset-specific observation. The sharper organization observed in the present analysis further motivated the family-based interpretation developed in the main manuscript.

**Evidence for harmonic families.** To determine whether the cross-subject similarity structure of individualized connectome harmonics reflects discrete higher-order organization rather than gradual similarity alone, we further examined the harmonic similarity matrix using hierarchical clustering (**Supplementary Figures 8–9**). Hierarchical clustering revealed a nested multiscale organization of the harmonic spectrum, from which a stable decomposition into harmonic families could be obtained. Importantly, harmonics assigned to the same family exhibited substantially greater mutual similarity than harmonics belonging to different families, independently validating the family organization introduced in the main manuscript. Together, these analyses indicate that the connectome harmonic spectrum possesses a reproducible higher-order organization that can be naturally described in terms of harmonic families, motivating the family-based analyses presented in the main manuscript.

**Structural organization of harmonic families.** Having established the existence of reproducible harmonic families, we next examined whether they preserve the multiscale structural organization observed for individual connectome harmonics (**Supplementary Figure 10**). Family-wise averages reproduced the same sequential evolution of graph support, sparsity, localization and spanned structural connectivity described in the main manuscript, indicating that the proposed family organization respects rather than obscures the intrinsic hierarchy of the harmonic spectrum. These findings further support the interpretation of harmonic families as higher-order structural units that retain the multiscale organization of individualized connectome harmonics.

**Robustness and generality of behavioral prediction.** To further evaluate the behavioral relevance of individualized connectome harmonics, we performed several complementary prediction analyses extending those presented in the main manuscript. These analyses assessed the specificity of the observed prediction profile across behavioral domains, its robustness to potential confounding by head movement, and the respective contributions of low- and high-frequency harmonics to behavioral prediction.

We first evaluated whether the cumulative prediction profile observed for cognition generalized to the remaining behavioral factors (**Supplementary Figure 11**). Unlike *Cognition*, neither *Mental health*, *Processing speed* nor *Substance use* exhibited consistent prediction above chance across the harmonic spectrum. These findings indicate that the multiscale structural organization identified in the present work preferentially captures inter-individual differences related to cognition.

To assess whether the principal behavioral findings could be explained by motion-related confounds, prediction analyses were repeated while including mean framewise displacement as a covariate of no interest (**Supplementary Figure 12**). The resulting prediction profile closely matched that obtained in the main manuscript, demonstrating that the observed relationships between connectome harmonics and cognition remain robust after accounting for head movement.

We examined how behavioral prediction evolved when structural connectivity was reconstructed by progressively incorporating harmonics from the highest rather than the lowest graph frequencies (**Supplementary Figure 13**). Although prediction again improved as additional harmonics were included, the accumulation profile differed markedly from that obtained when starting from low-frequency harmonics. Together, these analyses reinforce the conclusion that cognition is encoded through complementary structural information distributed across multiple spectral scales rather than being dominated by either end of the spectrum alone.

**Behavioral relevance of harmonic families.** Finally, we investigated whether the family organization preserves the behavioral information carried by individual connectome harmonics (**Supplementary Figure 14**). Accumulating structural connectivity reconstructions at the level of harmonic families reproduced the principal transitions observed for harmonic-wise prediction, despite representing the spectrum using a substantially smaller number of units. These findings suggest that harmonic families constitute a compact representation of the behavioral organization of the connectome harmonic spectrum, preserving its principal predictive transitions while substantially reducing its dimensionality.

*Figure 3. Sparse harmonic representations and temporal dynamics*

**Head movement across paradigms.** Motion statistics were first computed separately for $SES_1$ and $SES_2$ acquisitions and subsequently averaged within acquisition type, as no systematic differences between phase-encoding directions were observed (**Supplementary Figure 15A**; **Supplementary Tables 2–4**).

Overall motion levels were low. Only 121 participants (13.83%) exhibited more than 10% scrubbed frames across all acquisitions, while 44 (5.03%) and 13 (1.49%) exceeded the 20% and 30% thresholds, respectively.

Mean framewise displacement differed significantly across acquisitions (repeated-measures ANOVA: $F_{8,866} = 8.87$, $p < 0.001$, partial $\eta^2 = 0.008$). However, although 11 of 36 pairwise comparisons reached statistical significance after Bonferroni correction, all associated effect sizes remained small (maximum absolute Cohen's $d = 0.28$; **Supplementary Figure 15B**), indicating that head movement was broadly comparable across paradigms.

Framewise displacement was consistently negatively correlated with the *Cognition* factor score across paradigms ($R = -0.298 \pm 0.026$ and $R = -0.275 \pm 0.018$ before and after scrubbing, respectively), consistent with previous reports identifying distinct categories of movers during MRI acquisition[17]. Consequently, mean framewise displacement was included as a covariate in analyses performed on structural or non-scrubbed functional data, whereas mean framewise displacement computed after scrubbing was used for analyses based on scrubbed fMRI time series. Together, these observations indicate that the task-related differences in harmonic dynamics reported in the main manuscript are unlikely to reflect systematic differences in head movement across paradigms.

**Robustness of sparse harmonic representations.** Before characterizing the temporal dynamics of connectome harmonic recruitment, we evaluated the stability of the sparse representation across all HCP paradigms (**Supplementary Figures 16–17**). Sparse connectome harmonic representations consistently reconstructed regional BOLD activity with high accuracy throughout both resting-state and task acquisitions, indicating that the quality of the decomposition remained stable over time and across behavioral conditions. Furthermore, the proportion of harmonics simultaneously expressed remained remarkably constant across paradigms, suggesting that cognitive demands primarily modify which harmonics are recruited and how they are expressed, rather than the overall complexity of the harmonic representation.

**Robustness of harmonic expression dynamics.** To further evaluate the robustness of the temporal metrics introduced in Figure 3, harmonic expression intensity, entry rate and duration were additionally computed using all expression weights, only positive weights, and only negative weights (**Supplementary Figures 18–20**). Across all HCP paradigms, the characteristic spectral profiles remained highly similar irrespective of expression polarity, demonstrating that the reported organization does not arise from preferential recruitment of either positive or negative harmonic contributions. Together with the consistency observed across resting-state and task paradigms, these analyses indicate that the spectral organization of harmonic expression dynamics reflects a stable property of connectome harmonics rather than a polarity-specific effect.

**Associations of dynamic metrics with head movement.** Because head movement may influence estimates of functional activity, we additionally examined its relationship with each harmonic expression metric (**Supplementary Figure 21**). Across harmonics and behavioral paradigms, correlations between mean framewise displacement and harmonic expression intensity, entry rate, or duration remained uniformly weak. These findings further support the interpretation that the spectral organization reported in **Figure 3** primarily reflects neurophysiological rather than motion-related differences.

*Figure 4. Cross-paradigm harmonic dynamics*

**Robustness to head movement.** To determine whether the reported cross-paradigm differences were influenced by head movement, the statistical analyses were repeated using two-way ANOVA models that explicitly included mean framewise displacement and its interaction with behavioral paradigm (**Supplementary Figure 22**). Across harmonic expression intensity, entry rate and duration, the task factor remained the dominant source of variance, whereas the effects of head movement and the task-by-movement interaction were generally small. Consistent with this observation, accounting for head movement produced only minor changes in the resulting statistical profiles. Together, these analyses demonstrate that the principal cross-paradigm findings reported in **Figure 4A** are largely insensitive to motion-related effects.

**Detailed cross-paradigm comparisons.** **Supplementary Figure 23** presents the complete set of pairwise task contrasts underlying the summary visualization shown in **Figure 4A**. Whereas the main figure emphasizes the combinations of temporal metrics exhibiting significant differences, the signed contrast maps additionally indicate the direction of each effect. These analyses provide a comprehensive view of how harmonic expression dynamics differ between individual behavioral paradigms while remaining consistent with the overall organization described in the main manuscript.

**Reproducibility of cross-paradigm differences.** Additional analyses further demonstrated the robustness of the reported task-related effects (**Supplementary Figures 24–25**). Rest-to-task contrasts obtained using $REST_1$ and $REST_2$ exhibited highly similar statistical profiles across all three temporal metrics, indicating that the reported differences do not depend on the particular resting-state acquisition used for comparison. Likewise, repeating the analyses without explicitly accounting for head movement yielded highly similar patterns of significant harmonic differences. Together, these observations demonstrate that the principal cross-paradigm organization reported in **Figure 4A** is reproducible across independent resting-state acquisitions and robust to alternative statistical modeling choices.

**Statistical significance of prediction performance.** Permutation testing was performed for all prediction models presented in **Figure 4B** by repeatedly randomizing cognition factor scores and repeating the complete prediction pipeline (**Supplementary Figure 26**). In every case, the observed prediction performance exceeded the corresponding permutation-derived null distribution, confirming that all models performed significantly above chance. These analyses demonstrate that the reported prediction accuracies cannot be explained by random associations between harmonic descriptors and behavioral scores.

**Robustness of prediction performance to head movement.** Prediction analyses were repeated without including head movement as a covariate of no interest (**Supplementary Figure 27**). The resulting prediction accuracies closely matched those reported in **Figure 4B**, with the same ordering of structural, functional and hybrid kernel models, and similar differences between matched-pair and all-pair representations. Together, these analyses indicate that the principal prediction findings are largely insensitive to the explicit modeling of head movement.

**Comparison with conventional connectivity measures.** To place the proposed harmonic-based representations in context, prediction performance was additionally evaluated using kernels constructed directly from structural connectivity or functional connectivity (**Supplementary Figure 28**). Functional connectivity achieved the highest predictive performance, followed by the proposed hybrid harmonic kernel and structural connectivity. These results indicate that the harmonic representation preserves substantial behaviorally relevant information while providing a compact joint description of structural and functional organization.

### Supplementary Tables

**Supplementary Table 1: Demographic characteristics of the study cohort.** Continuous variables are reported as mean ± standard deviation, while categorical variables are reported as counts.

| **Characteristic** | **Value** |
|---|---|
| Number of participants | 875 |
| Age (years) | $28.67 \pm 3.70$ |
| Sex (male / female) | 412 / 463 |
| *Ethnicity* | |
| Hispanic or Latino | 81 |
| Not Hispanic or Latino | 794 |
| *Race* | |
| White | 660 |
| Black or African American | 121 |
| Asian, Hawaiian or Other Pacific Islands | 56 |
| American Indian/Alaskan Native | 1 |
| More Than One | 21 |
| Unknown | 17 |

**Supplementary Table 2: Summary of mean framewise displacement across functional MRI acquisitions.** Mean framewise displacement in mm was computed for each participant and summarized separately for each acquisition. The 95% interval corresponds to the 2.5$^{\text{th}}$–97.5$^{\text{th}}$ percentiles.

| **Acquisition** | **Mean** | **Median** | **95% interval** | **Min** | **Max** |
|---|---|---|---|---|---|
| EMOTION | 0.158 | 0.147 | 0.083–0.306 | 0.061 | 0.523 |
| GAMBLING | 0.156 | 0.143 | 0.086–0.298 | 0.07 | 0.826 |
| LANGUAGE | 0.164 | 0.151 | 0.089–0.316 | 0.071 | 0.507 |
| MOTOR | 0.172 | 0.16 | 0.098–0.315 | 0.077 | 0.734 |
| RELATIONAL | 0.168 | 0.154 | 0.088–0.339 | 0.056 | 0.664 |
| SOCIAL | 0.16 | 0.148 | 0.084–0.306 | 0.061 | 0.417 |
| WM | 0.16 | 0.147 | 0.088–0.286 | 0.066 | 0.648 |
| $\text{REST}_1$ | 0.158 | 0.146 | 0.089–0.303 | 0.072 | 0.531 |
| $\text{REST}_2$ | 0.168 | 0.151 | 0.091–0.34 | 0.068 | 0.534 |

**Supplementary Table 3: Summary of mean framewise displacement across functional MRI acquisitions (non-scrubbed frames).** Mean framewise displacement in mm across non-scrubbed frames at a threshold of 0.5 mm[18] was computed for each participant and summarized separately for each acquisition. The 95% interval corresponds to the 2.5$^{th}$–97.5$^{th}$ percentiles.

| **Acquisition** | **Mean** | **Median** | **95% interval** | **Min** | **Max** |
|---|---|---|---|---|---|
| EMOTION | 0.151 | 0.144 | 0.082–0.264 | 0.061 | 0.309 |
| GAMBLING | 0.15 | 0.14 | 0.084–0.265 | 0.07 | 0.313 |
| LANGUAGE | 0.156 | 0.147 | 0.088–0.269 | 0.07 | 0.296 |
| MOTOR | 0.159 | 0.152 | 0.097–0.26 | 0.077 | 0.309 |
| RELATIONAL | 0.157 | 0.149 | 0.087–0.268 | 0.056 | 0.304 |
| SOCIAL | 0.151 | 0.143 | 0.083–0.259 | 0.061 | 0.305 |
| WM | 0.152 | 0.144 | 0.087–0.262 | 0.066 | 0.31 |
| $REST_1$ | 0.152 | 0.144 | 0.089–0.265 | 0.069 | 0.309 |
| $REST_2$ | 0.157 | 0.148 | 0.09–0.27 | 0.068 | 0.308 |

**Supplementary Table 4: Summary of percentage of scrubbed frames across functional MRI acquisitions.** Percentage of scrubbed frames at a framewise displacement threshold of 0.5 mm[18] was computed for each participant and summarized separately for each acquisition. The 95% interval corresponds to the $2.5^{th}$–$97.5^{th}$ percentiles.

| Acquisition | Mean | Median | 95% interval | Min | Max |
|---|---|---|---|---|---|
| EMOTION | 1.23 | 0 | 0–10 | 0 | 43.529 |
| GAMBLING | 0.984 | 0 | 0–7.945 | 0 | 30.769 |
| LANGUAGE | 1.315 | 0 | 0–11.371 | 0 | 36.774 |
| MOTOR | 2.273 | 0.719 | 0–12.68 | 0 | 35.252 |
| RELATIONAL | 1.789 | 0.443 | 0–15.597 | 0 | 34.071 |
| SOCIAL | 1.422 | 0 | 0–12.174 | 0 | 37.687 |
| WM | 1.237 | 0.251 | 0–8.02 | 0 | 39.599 |
| $REST_1$ | 1.286 | 0.168 | 0–11.087 | 0 | 49.246 |
| $REST_2$ | 2.04 | 0.335 | 0–17.379 | 0 | 54.523 |

### Supplementary Figures

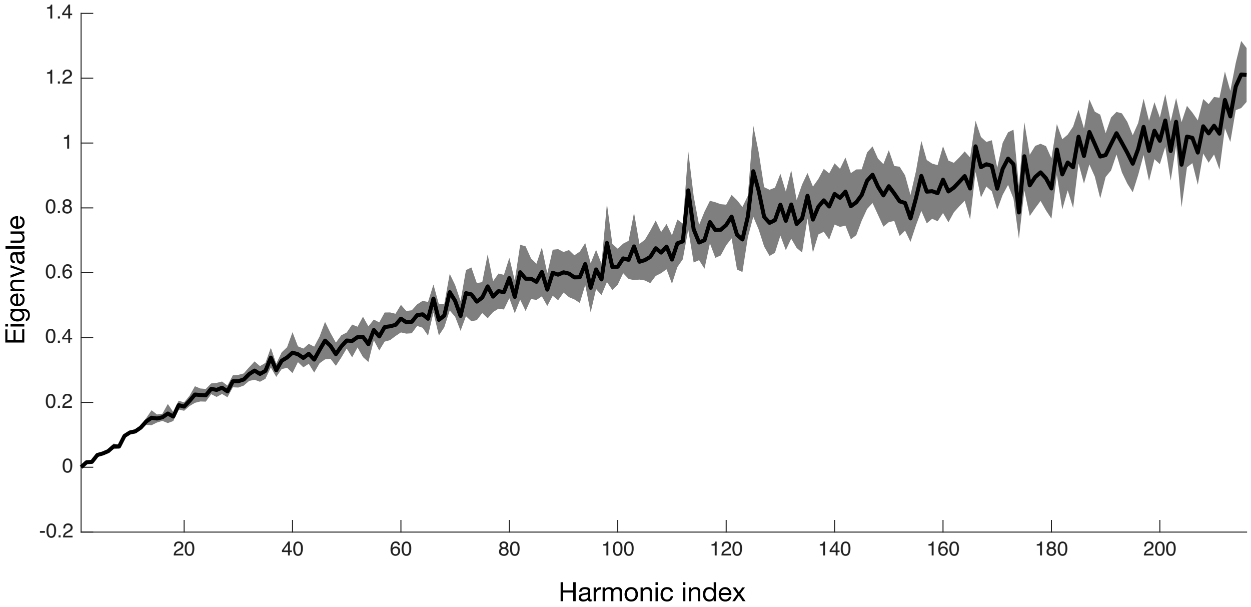

**Supplementary Figure 1: Connectome harmonic eigenvalues increase smoothly with harmonic index.** Eigenvalues of matched connectome harmonics as a function of harmonic index. Shaded regions denote standard deviation across subjects. The smooth increase confirms that the harmonic matching procedure preserves the spectral ordering of connectome harmonics across individuals.

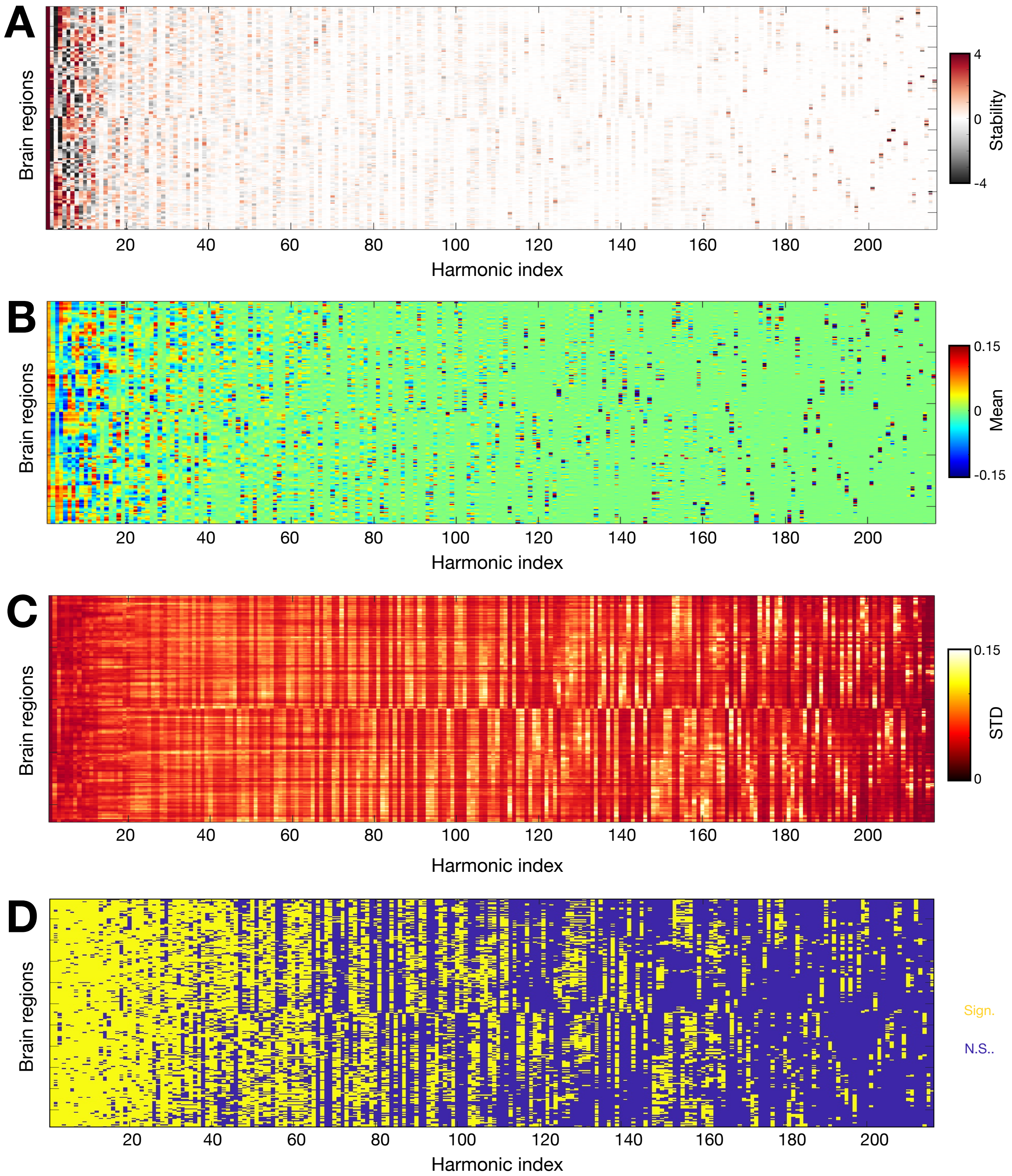

**Supplementary Figure 2: Regional instability of intermediate-frequency harmonics reflects increased inter-subject variability rather than loss of consistent signal.** **(A)** Population-wise regional stability (reproduced from **Figure 1B**). **(B)** Population-wise mean of harmonic signal across subjects. **(C)** Population-wise standard deviation of harmonic signal across subjects. **(D)** Brain region–harmonic combinations exhibiting significant stability after Bonferroni correction are shown in yellow. Together, these analyses show that although stability decreases for intermediate-frequency harmonics, many regional harmonic patterns remain statistically reproducible across subjects. STD: standard deviation.

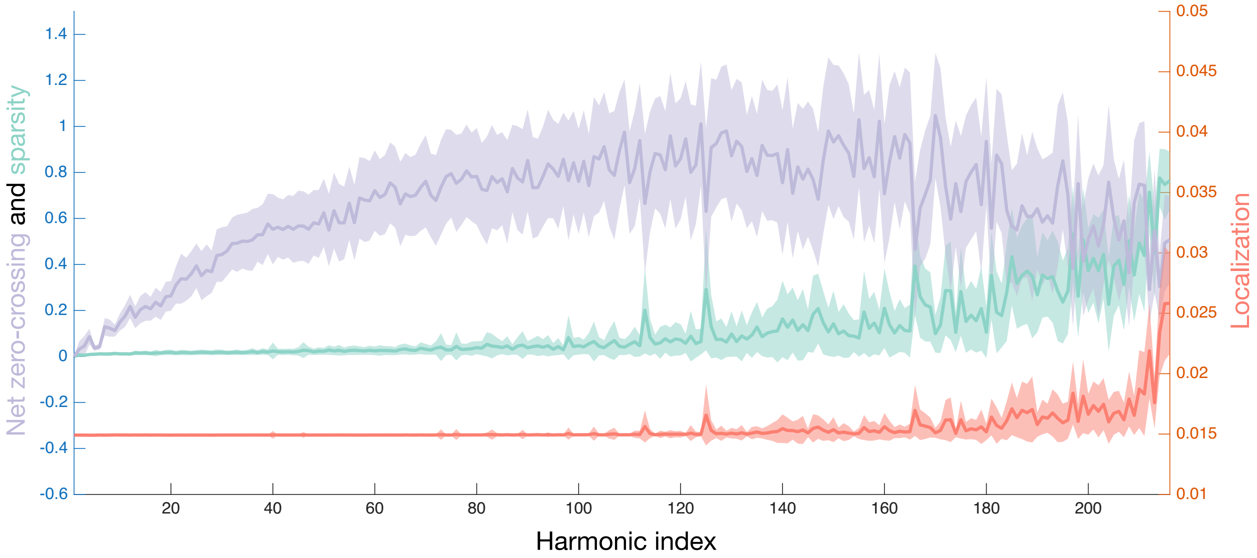

**Supplementary Figure 3: Structural descriptors evolve coherently across the harmonic spectrum.** Net zero-crossing (light purple), computed as in Sipes et al.[15]; harmonic sparsity (turquoise), defined as the fraction of supra-threshold brain regions; and localization (salmon), defined as the mean inverse Euclidean distance between pairs of supra-threshold brain regions, shown as a function of harmonic index. Shaded regions denote standard deviation across subjects. These results reproduce previous observations by Sipes et al.[15] and further show that increasing harmonic sparsity is accompanied by progressively localized structural representations.

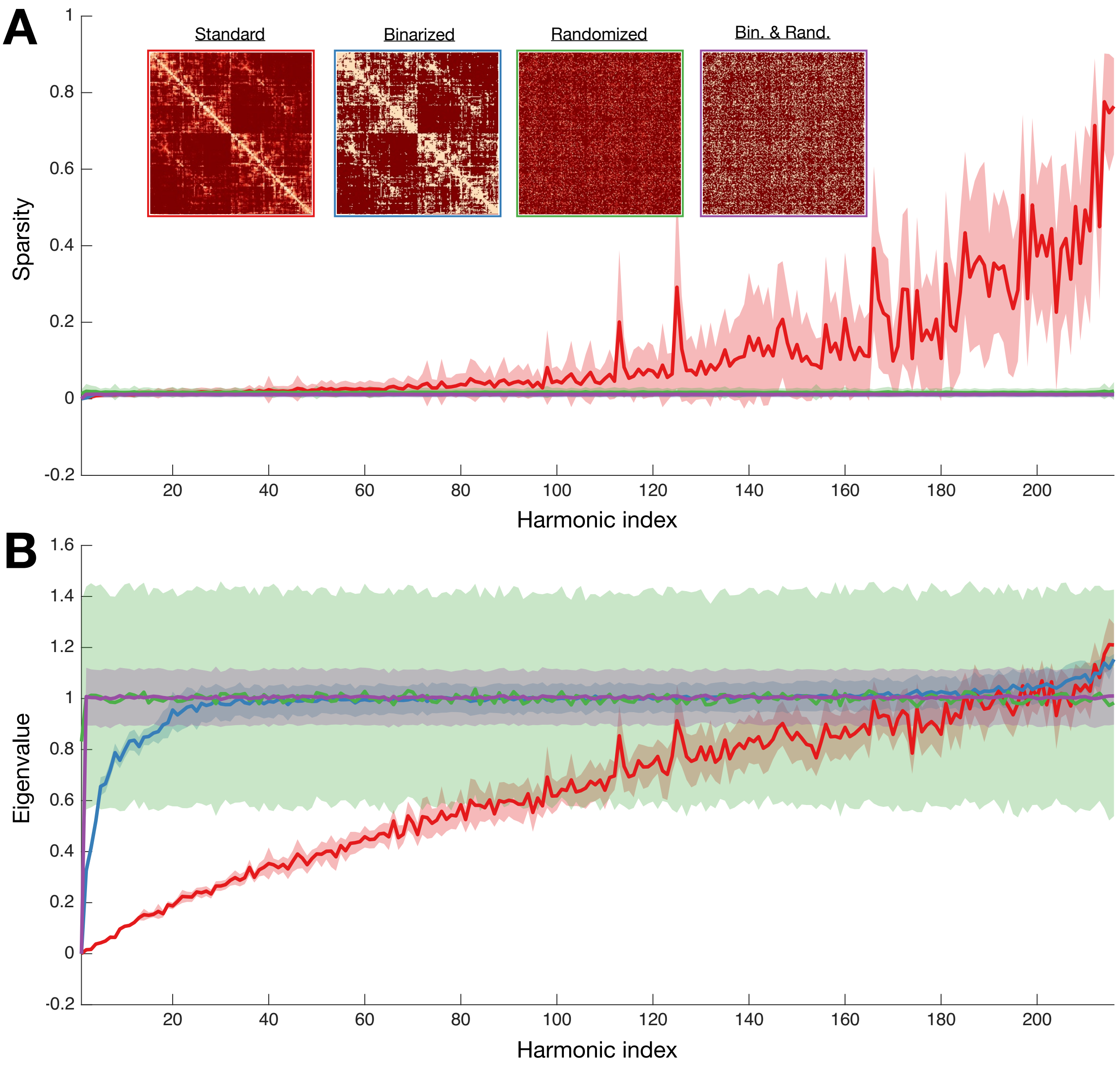

**Supplementary Figure 4: Connectivity weights and graph topology jointly determine the emergence of harmonic sparsity.** **(A)** Harmonic sparsity (fraction of supra-threshold brain regions) as a function of harmonic index for connectome harmonics extracted from standard structural connectivity matrices (red), binarized matrices (blue), degree-preserving randomized matrices (green), or matrices subjected to both manipulations (purple). (*Inset*) Representative structural connectivity matrices for each condition. **(B)** Harmonic eigenvalues for the same four connectivity conditions. Shaded regions denote standard deviation across subjects. Together, these analyses demonstrate that both weighted connectivity and native graph topology are required for the emergence of harmonic sparsity. Bin.: binarized, Rand.: randomized.

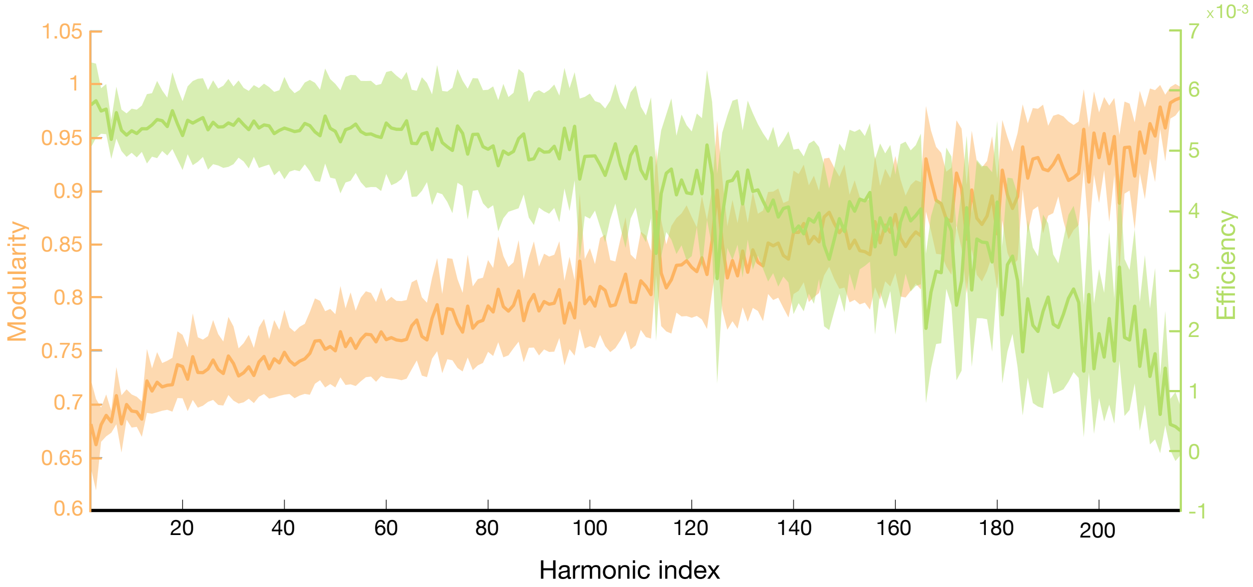

**Supplementary Figure 5: Graph-theoretical properties of harmonic-wise structural connectivity evolve smoothly across the graph frequency spectrum.** Modularity (orange) and global efficiency (green) as a function of harmonic index. Graph-theoretical measures were computed on structural connectivity matrices reconstructed from individual connectome harmonics. Shaded regions denote standard deviation across subjects. Together, these analyses show that harmonic-specific structural connectivity gradually transitions from globally integrated to increasingly modular organization across the spectrum.

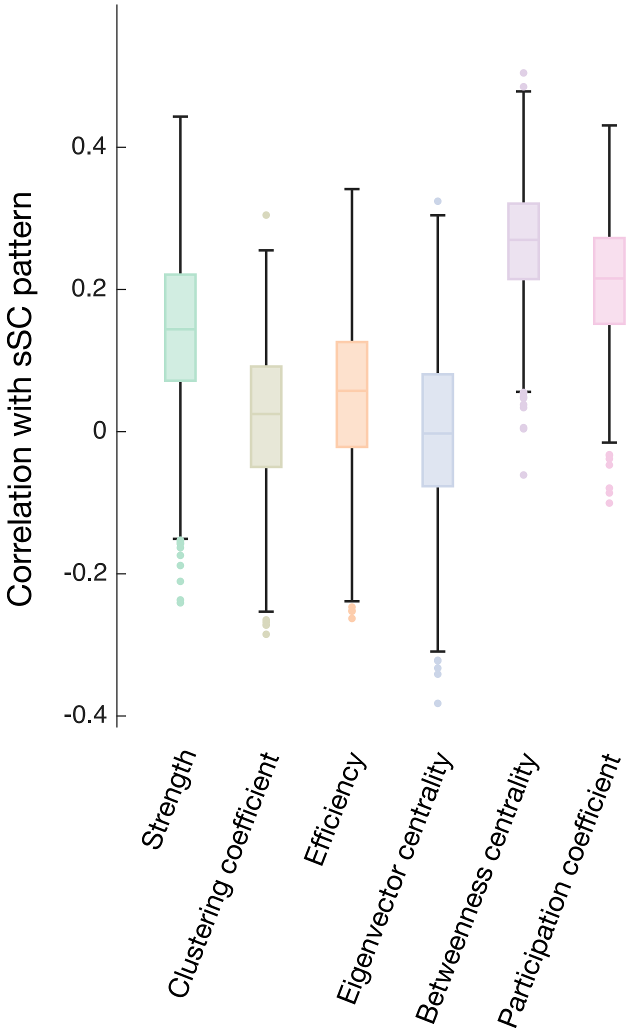

**Supplementary Figure 6: Spectral participation is most strongly associated with graph-theoretical measures of network integration.** Regional strength, clustering coefficient, global efficiency, eigenvector centrality, betweenness centrality and participation coefficient were computed from each subject's structural connectivity matrix. For each graph-theoretical measure, its regional profile was correlated with the corresponding profile of spectral participation, defined as the number of connectome harmonics for which a brain region exhibited supra-threshold signal. Box plots summarize the distribution of correlation coefficients across subjects. Given the large sample size, all correlations were significantly greater than zero ($p < 0.001$). Together, these analyses indicate that spectral participation primarily reflects the integrative role of brain regions within the structural connectome.

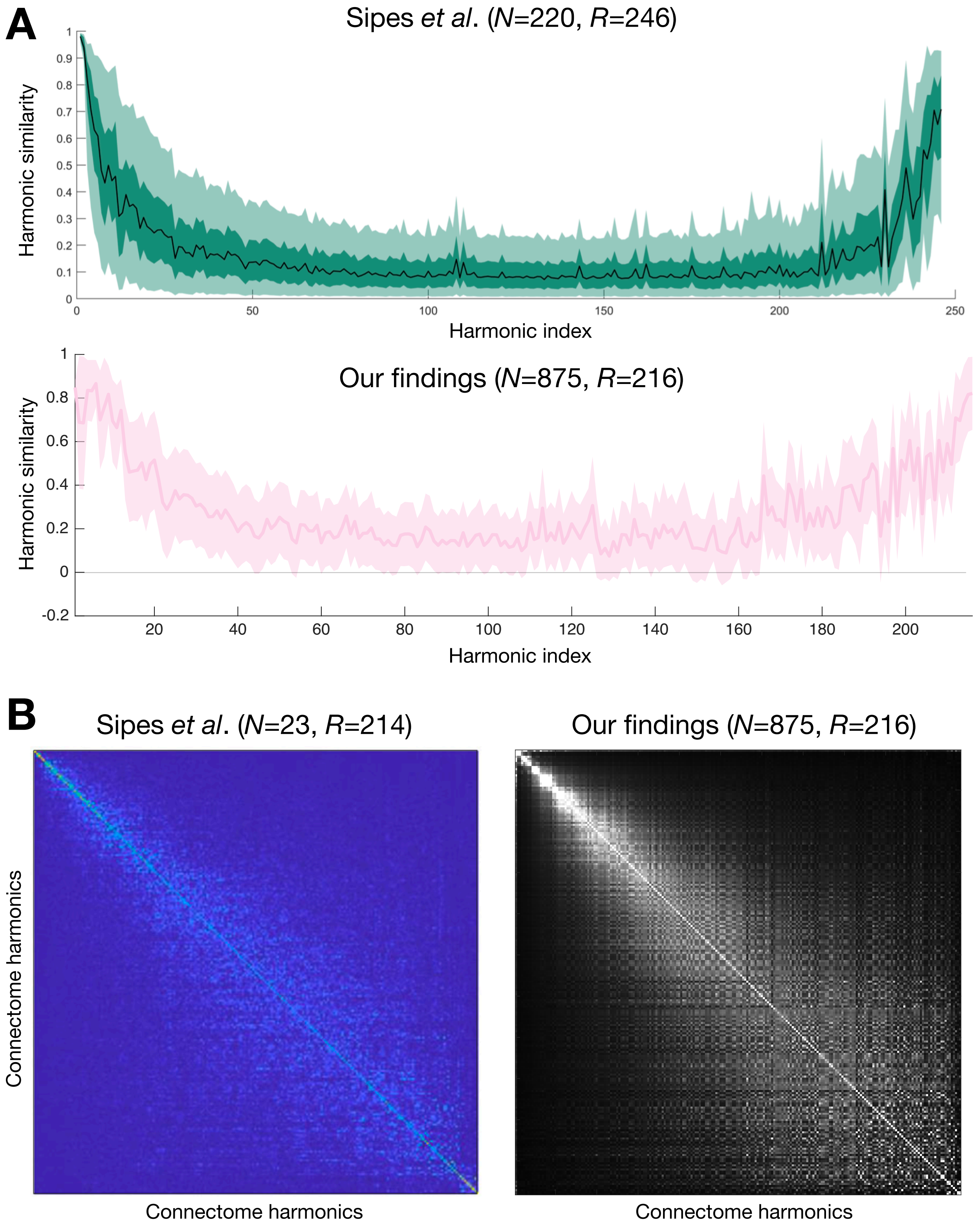

**Supplementary Figure 7: Reproduction of previously reported cross-subject similarity patterns. (A)** Similarity between matched connectome harmonics as a function of harmonic index, reproduced from Sipes et al.[15] (top) and obtained in the present work (bottom). Despite differences in dataset and methodological choices, both analyses reveal the same characteristic U-shaped relationship between harmonic similarity and graph frequency. **(B)** Pairwise similarity between all connectome harmonics in Sipes et al.[15] (left) and in the present work (right). The overall organization is qualitatively similar in both studies, although the block structure appears more clearly in the present analysis owing to the use of absolute dot products as similarity measure.

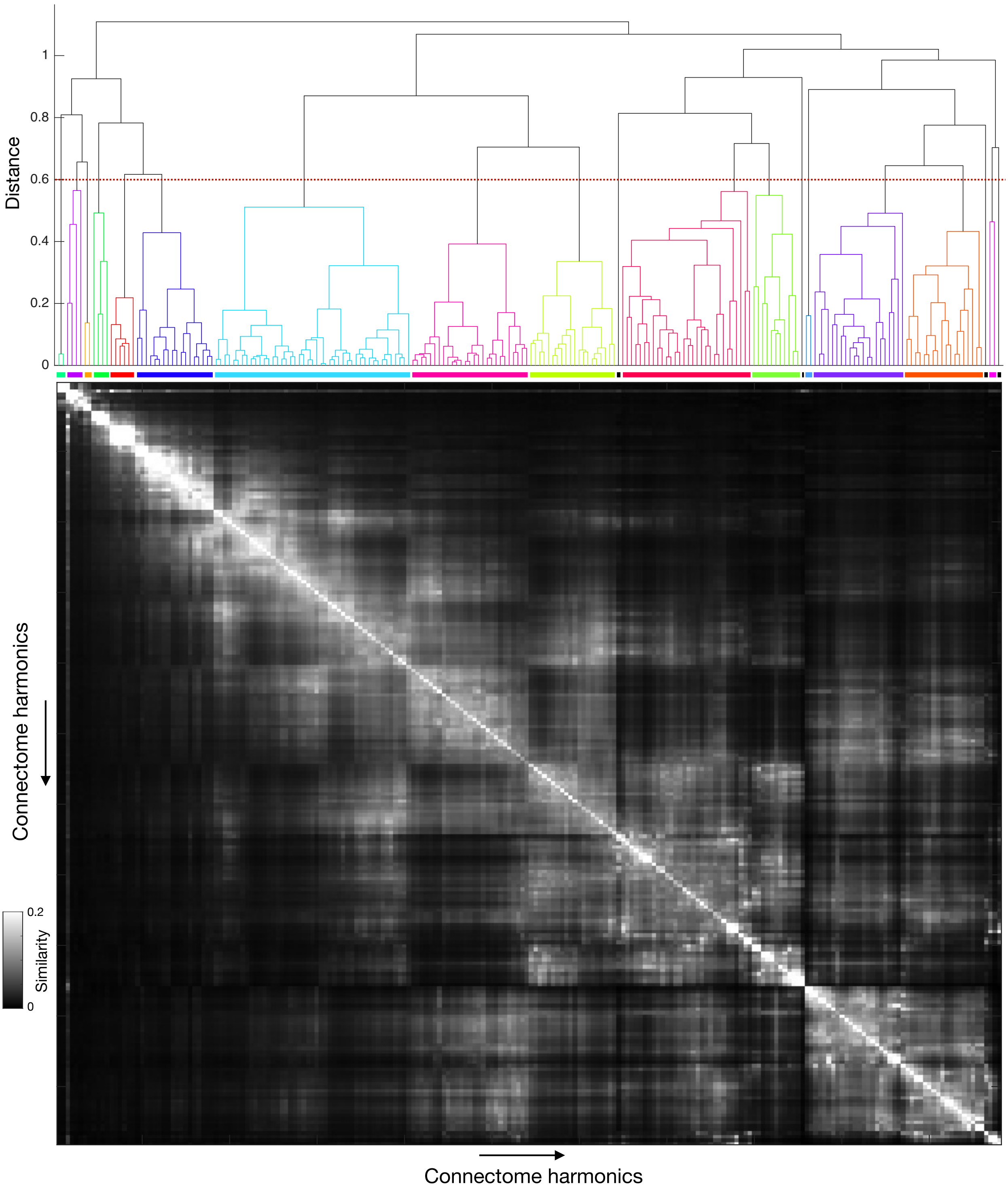

**Supplementary Figure 8: Hierarchical clustering reveals a multiscale family organization of connectome harmonics.** Pairwise similarity matrix reordered according to hierarchical clustering of harmonic-wise similarity profiles. The dendrogram (top) summarizes the resulting multiscale organization. Cutting the dendrogram at a clustering distance of 0.6 (red dashed line) yielded 19 harmonic families while preserving the prominent low-frequency similarity blocks shown in **Figure 2A**. Together, these analyses demonstrate that harmonic similarity exhibits a nested multiscale organization, supporting the interpretation of connectome harmonics as reproducible spectral families rather than isolated eigenmodes.

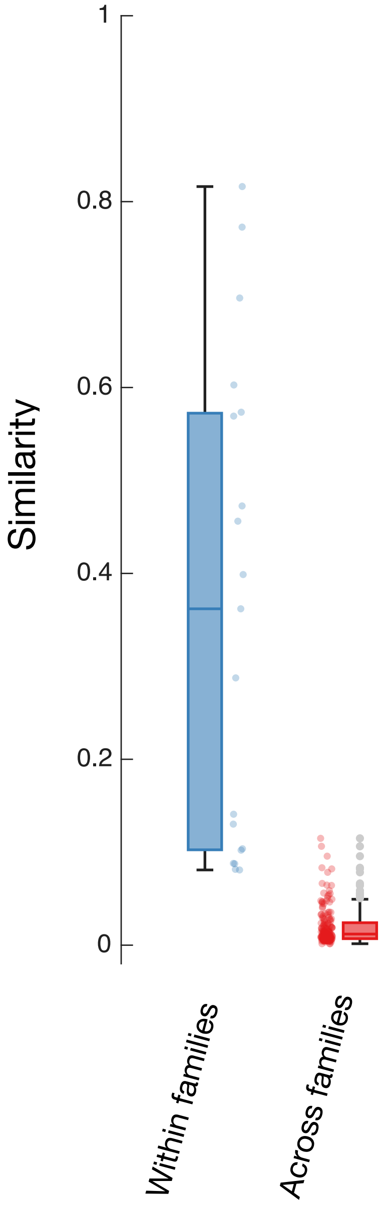

**Supplementary Figure 9: Within-family similarity largely exceeds cross-family similarity.** Average similarity within individual harmonic families (blue) and between pairs of distinct families (red). Box plots summarize family-wise mean similarity values. Both distributions differed significantly ($p < 10^{-6}$). These observations validate the family decomposition used throughout the manuscript.

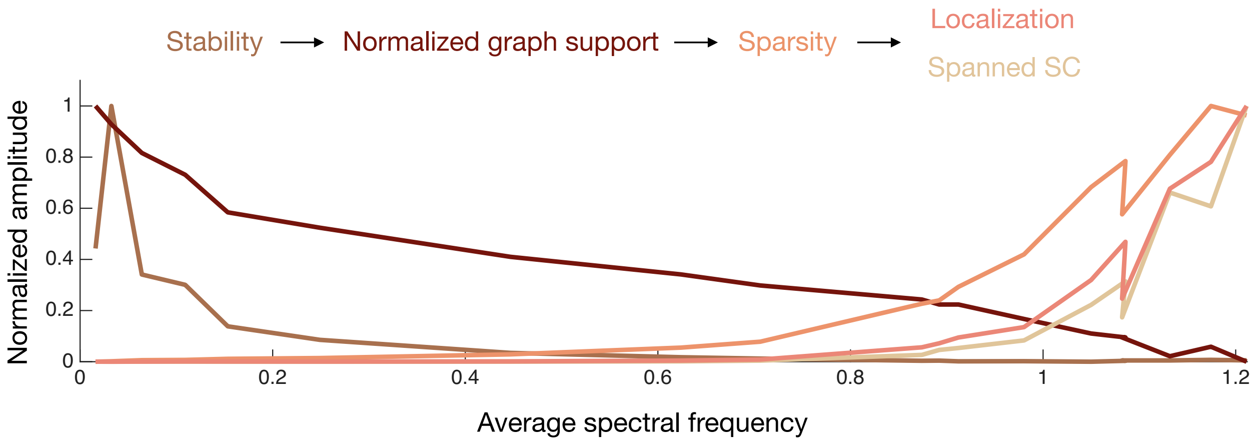

**Supplementary Figure 10: Harmonic families preserve the structural progression observed at the individual harmonic level.** Population-average family-wise stability (brown), normalized graph support (dark brown), sparsity (orange), localization (salmon) and spanned structural connectivity (light brown) are shown as a function of mean family spectral frequency. Subject-specific values were first averaged across subjects within each family. Each descriptor was then linearly normalized between 0 and 1 to facilitate comparison of their relative evolution across the spectrum. Together, these analyses demonstrate that the family-wise organization preserves the same sequential structural specialization observed for individual harmonics, with changes in stability preceding reductions in graph support, followed by increases in sparsity, anatomical localization and spanned structural connectivity.

P1_SF11_Cropped.pdf

**Supplementary Figure 11: Prediction of *Mental health*, *Processing speed* and *Substance use*.** Coefficient of determination $R^2$ obtained on left-out data, in a nested cross-validation scheme, when predicting the *Mental health* **(A)**, *Processing speed* **(B)** and *Substance use* **(C)** factor scores using structural connectivity approximations reconstructed with cumulatively increasing numbers of harmonics (X-axis), starting from the lowest-frequency ones. Shaded regions denote standard error of the mean. These results show that successful prediction is specific to the *Cognition* factor score.

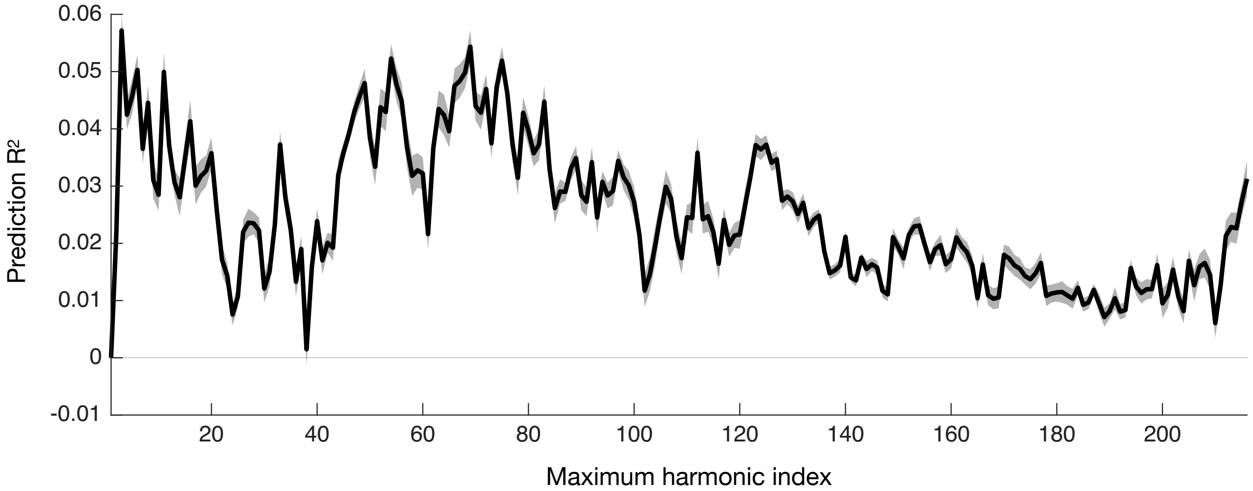

**Supplementary Figure 12: Prediction of cognition is insensitive to head movement.** Coefficient of determination $R^2$ obtained on left-out data, in a nested cross-validation scheme, when predicting the *Cognition* factor score using structural connectivity approximations reconstructed with cumulatively increasing numbers of harmonics (X-axis), starting from the lowest-frequency ones, and including mean framewise displacement as a covariate of no interest. Shaded regions denote standard error of the mean. The spectral profile closely resembles that shown in **Figure 2B**, with prediction accuracy consistently shifted upward by approximately 0.02. Together, these analyses demonstrate that the principal prediction results are insensitive to accounting for head movement.

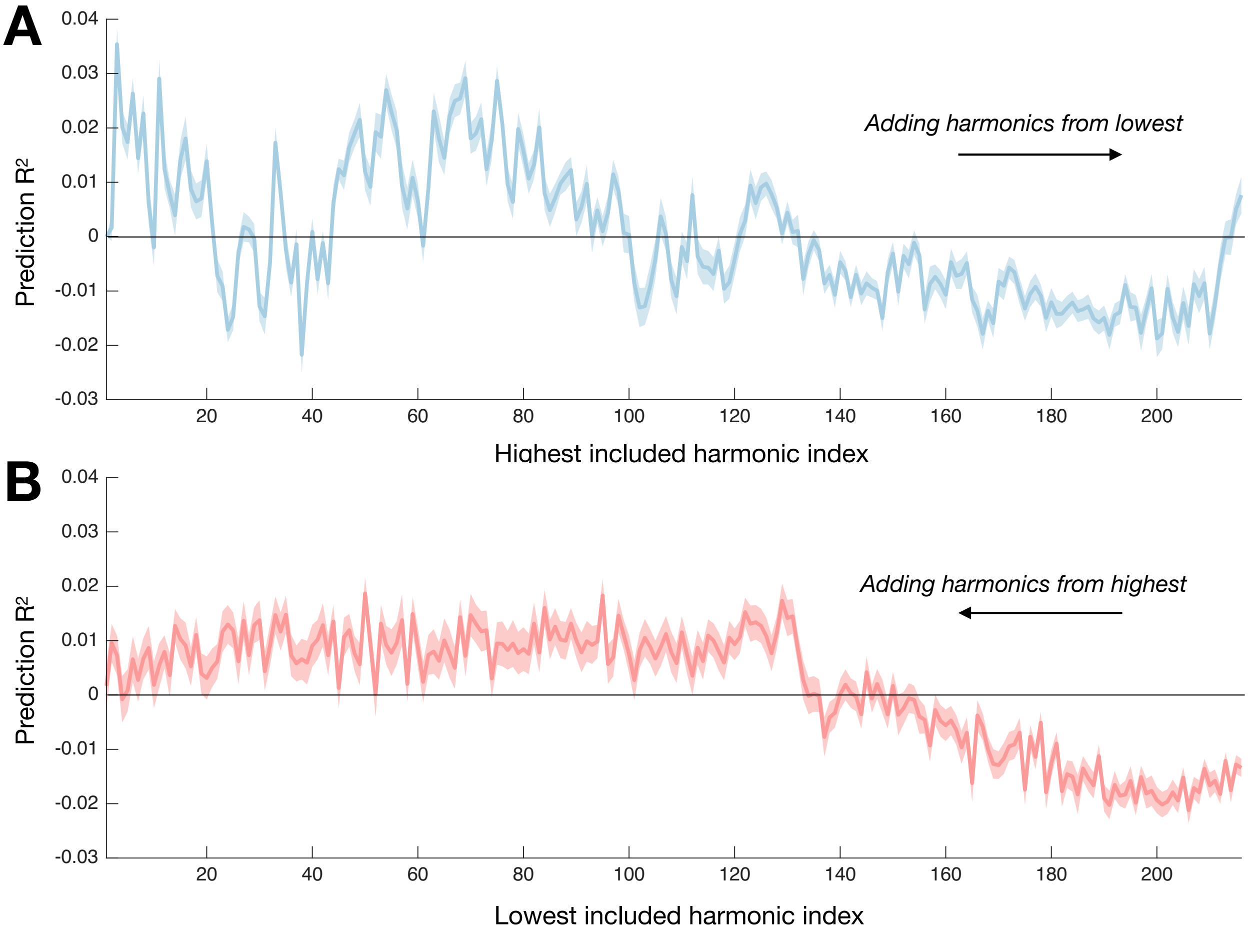

**Supplementary Figure 13: Prediction profiles differ when harmonics are accumulated from opposite ends of the spectrum.** Coefficient of determination $R^2$ obtained on left-out data in a nested cross-validation scheme when predicting the *Cognition* factor score using structural connectivity approximations reconstructed by cumulatively adding harmonics from the lowest frequencies upward **(A)** or from the highest frequencies downward **(B)**. Shaded regions denote standard error of the mean. The distinct accumulation trajectories demonstrate that cognition is encoded through complementary contributions from multiple spectral ranges rather than by either low- or high-frequency harmonics alone.

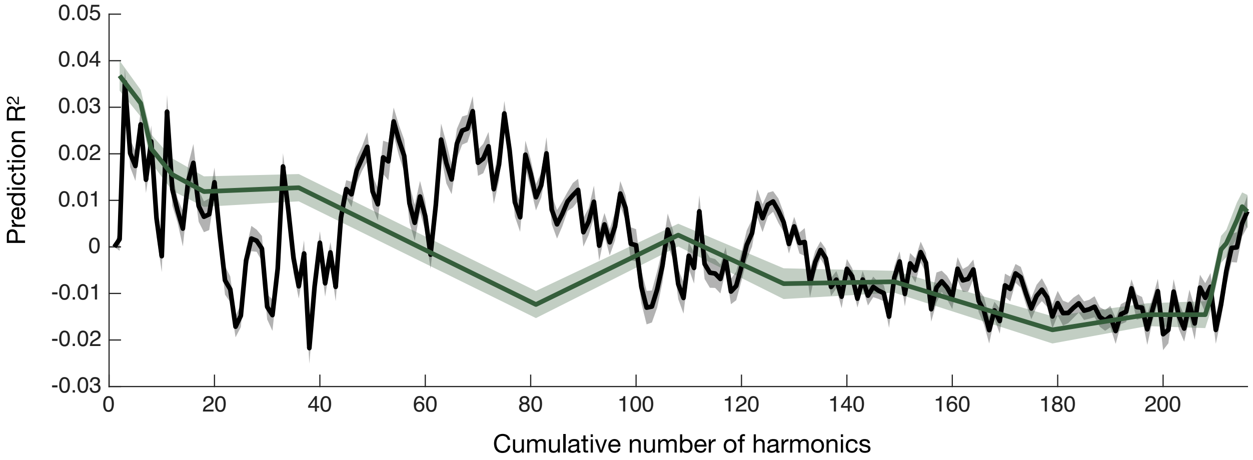

**Supplementary Figure 14: Harmonic families capture the major transitions in cumulative prediction performance.** Coefficient of determination $R^2$ obtained on left-out data in a nested cross-validation scheme when predicting the *Cognition* factor score using structural connectivity approximations reconstructed by cumulatively adding harmonics from the lowest frequencies upward (black), or using structural connectivity approximations reconstructed by cumulatively adding individual harmonics (black) or harmonic families (green), ordered by increasing graph frequency. The family-wise accumulation profile closely follows the large-scale evolution of the harmonic-wise prediction curve, indicating that harmonic families preserve the principal transitions in predictive information while providing a substantially more compact representation.

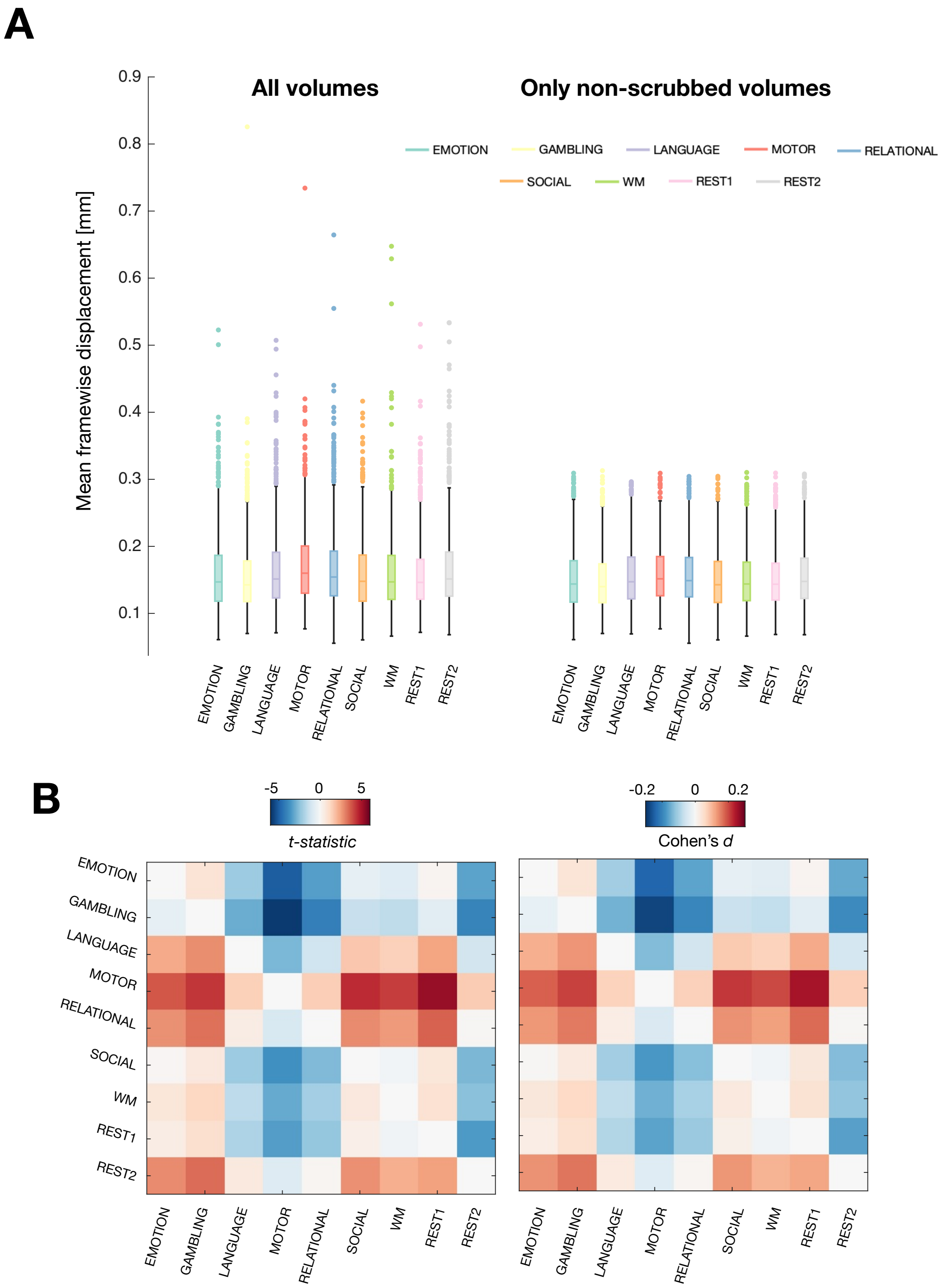

**Supplementary Figure 15: Head movement is broadly comparable across HCP paradigms. (A)** Mean framewise displacement computed from all fMRI volumes (left) or only from volumes retained after scrubbing at 0.5 mm[12] (right), shown separately for each HCP paradigm. **(B)** Pairwise task comparisons of mean framewise displacement expressed as *t*-statistics (left) and effect sizes (Cohen's *d*; right). Upper triangular entries correspond to analyses using all volumes, whereas lower triangular entries correspond to analyses after scrubbing. Together, these analyses indicate that differences in head movement between HCP paradigms were generally small and were further reduced after scrubbing, supporting the interpretation that the task-related effects reported in **Figure 4** primarily reflect differences in harmonic dynamics rather than systematic motion differences.

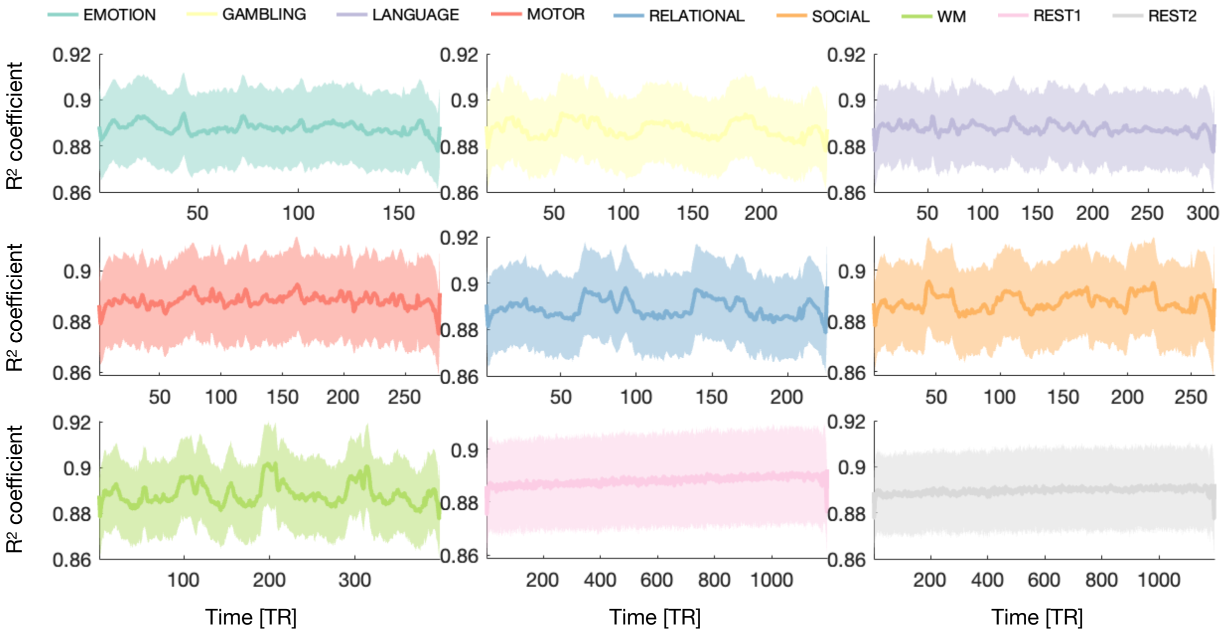

**Supplementary Figure 16: Sparse connectome harmonic representations accurately reconstruct regional activity across all HCP paradigms.** For all paradigms from the HCP dataset, coefficient of determination $R^2$ between observed and reconstructed regional BOLD activity shown as a function of time for each HCP paradigm. Shaded regions denote standard deviation across subjects. These results demonstrate that sparse connectome harmonic representations provide accurate and temporally stable reconstructions of regional brain activity across both resting-state and task paradigms.

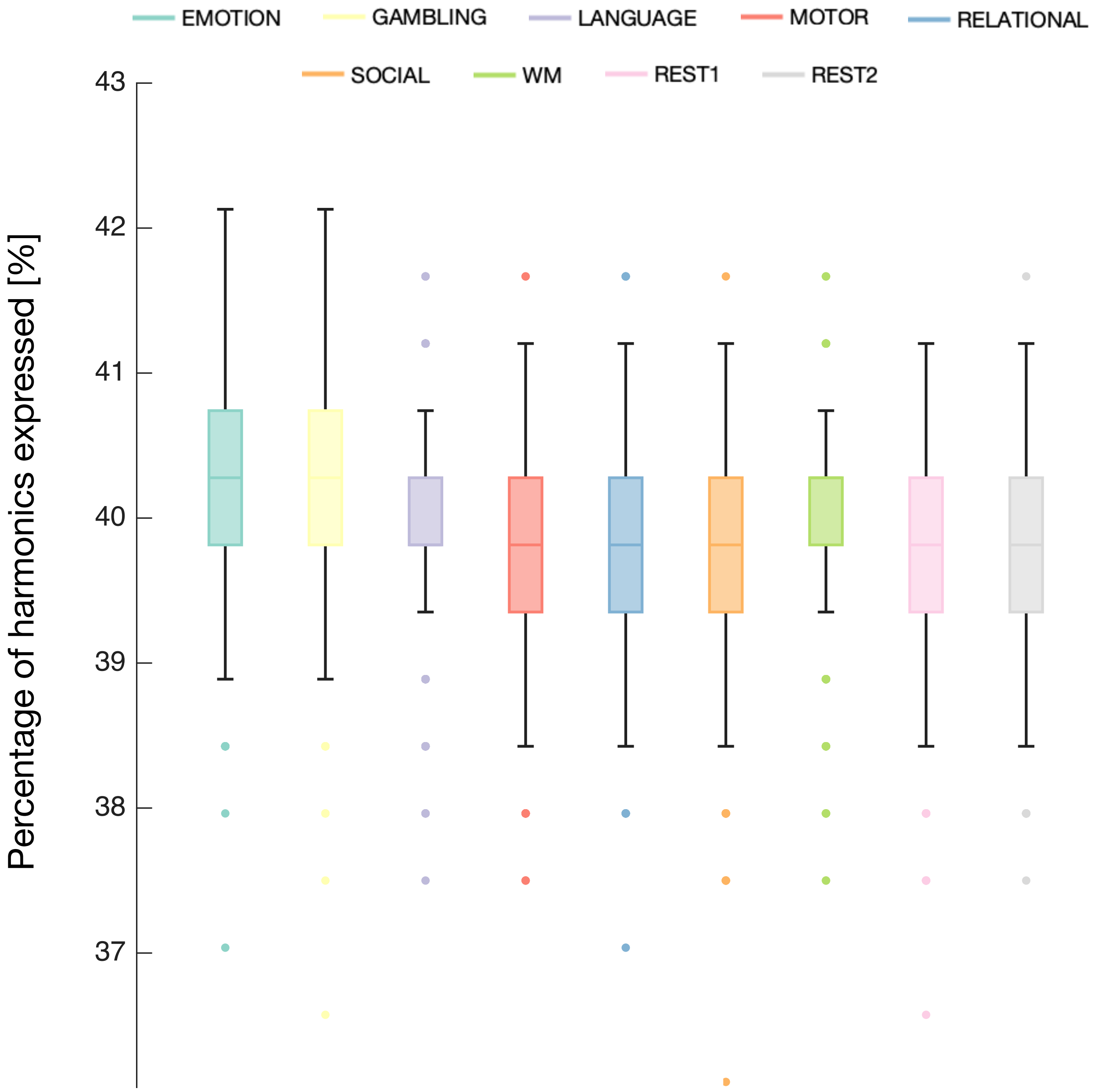

**Supplementary Figure 17: The sparsity of connectome harmonic expression is preserved across behavioral paradigms.** Distribution across subjects of the mean percentage of connectome harmonics simultaneously expressed over time for each HCP paradigm. The remarkably consistent sparsity across resting-state and task paradigms indicates that behavioral state primarily alters the identity and temporal dynamics of recruited harmonics rather than the overall fraction of the harmonic spectrum expressed at any given time.

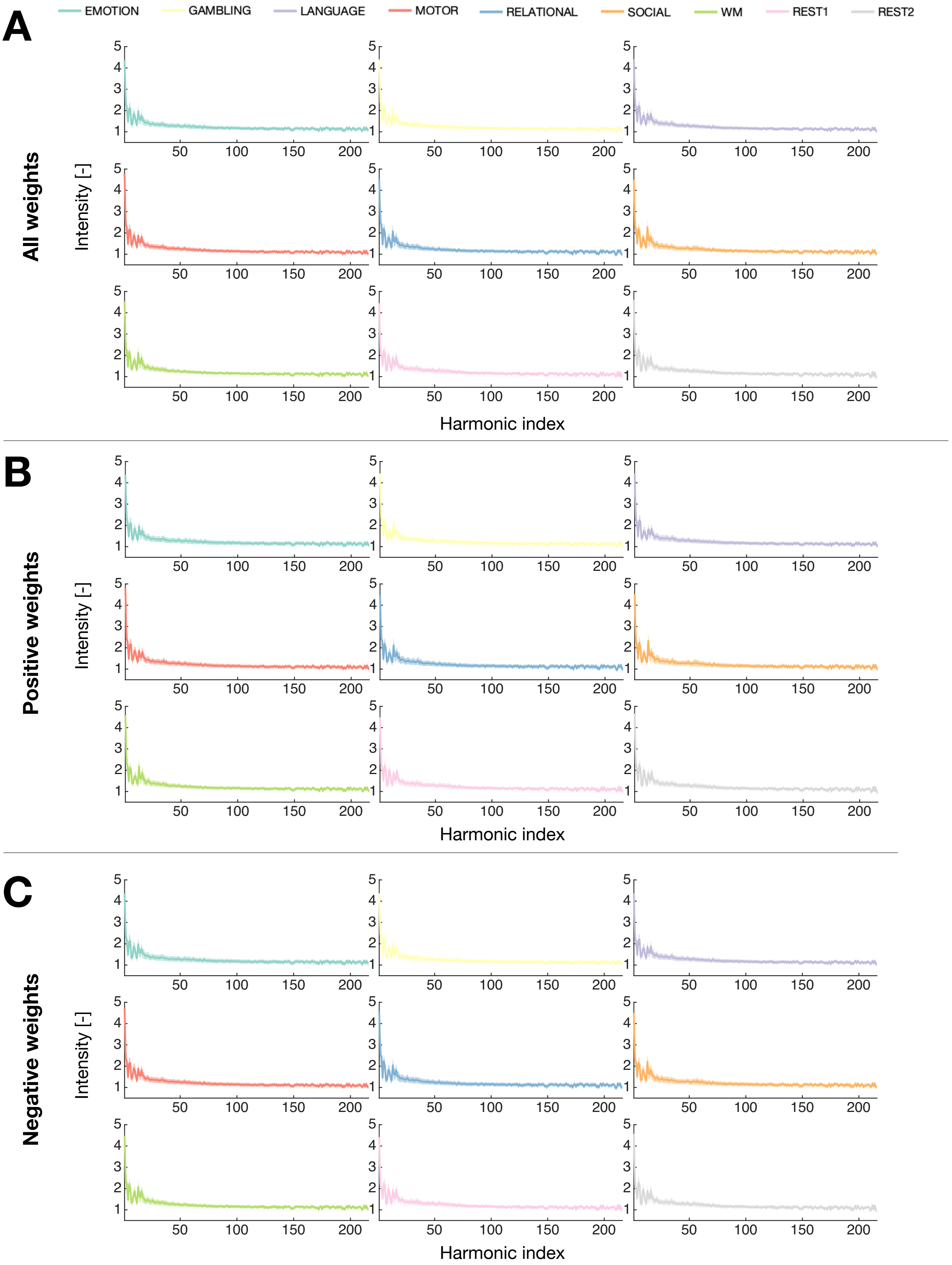

**Supplementary Figure 18: Spectral profiles of harmonic expression intensity are robust across paradigms and expression polarity.** Mean harmonic expression intensity as a function of harmonic index for each HCP paradigm, computed using **(A)** all expression weights, **(B)** only positive weights, or **(C)** only negative weights. Shaded regions denote standard deviation across subjects. The spectral organization of harmonic expression intensity is highly consistent across behavioral paradigms and remains highly similar when positive and negative expression weights are analyzed separately, indicating that the observed dynamics are not driven by a single expression polarity.

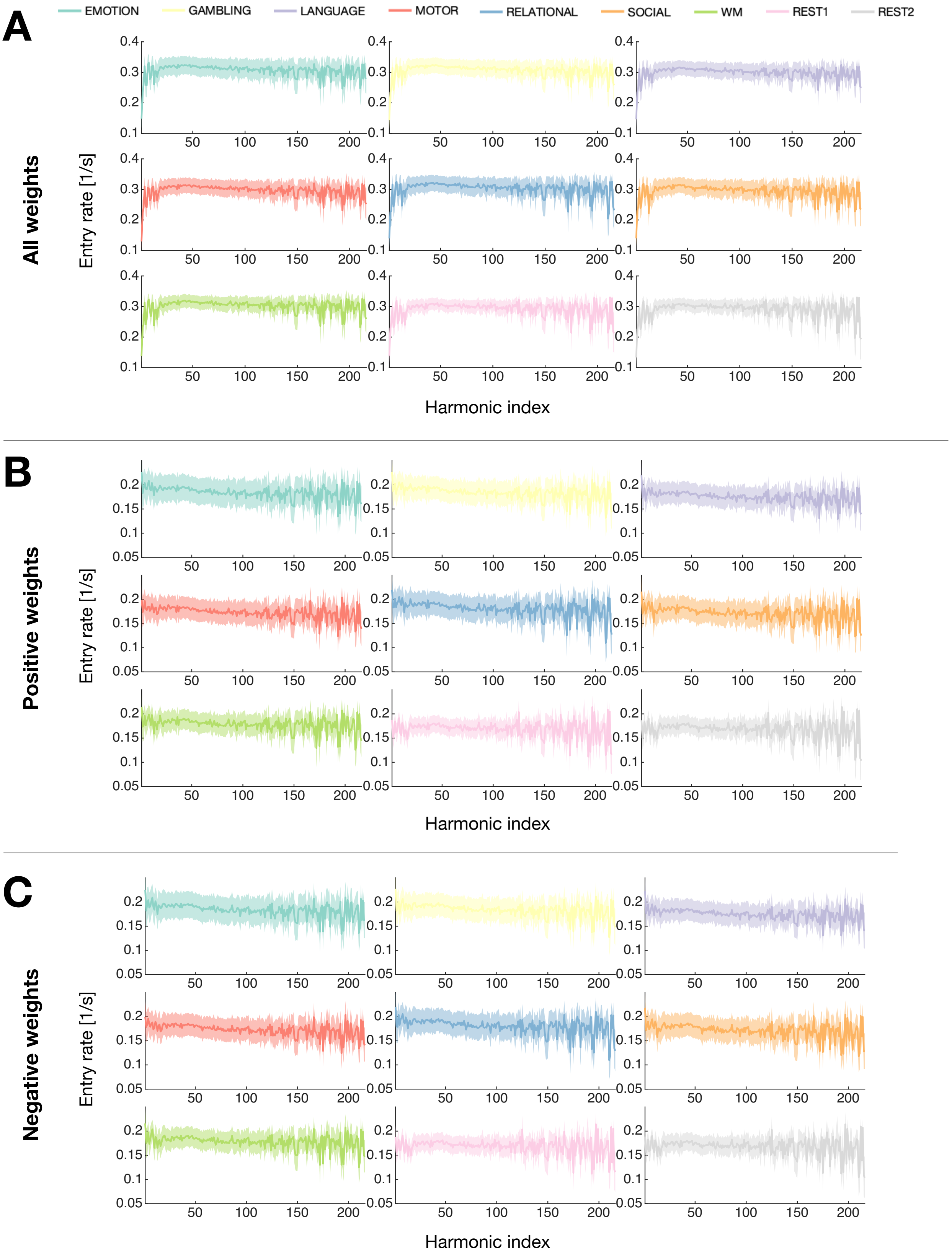

**Supplementary Figure 19: Spectral profiles of harmonic expression entry rate are preserved across paradigms and expression polarity.** Mean harmonic expression entry rate as a function of harmonic index for each HCP paradigm, computed using **(A)** all expression weights, **(B)** only positive weights, or **(C)** only negative weights. Shaded regions denote standard deviation across subjects. The spectral organization of harmonic expression entry rate is highly consistent across paradigms and remains highly similar when positive and negative expression weights are analyzed separately, indicating that the observed dynamics are not driven by a single expression polarity.

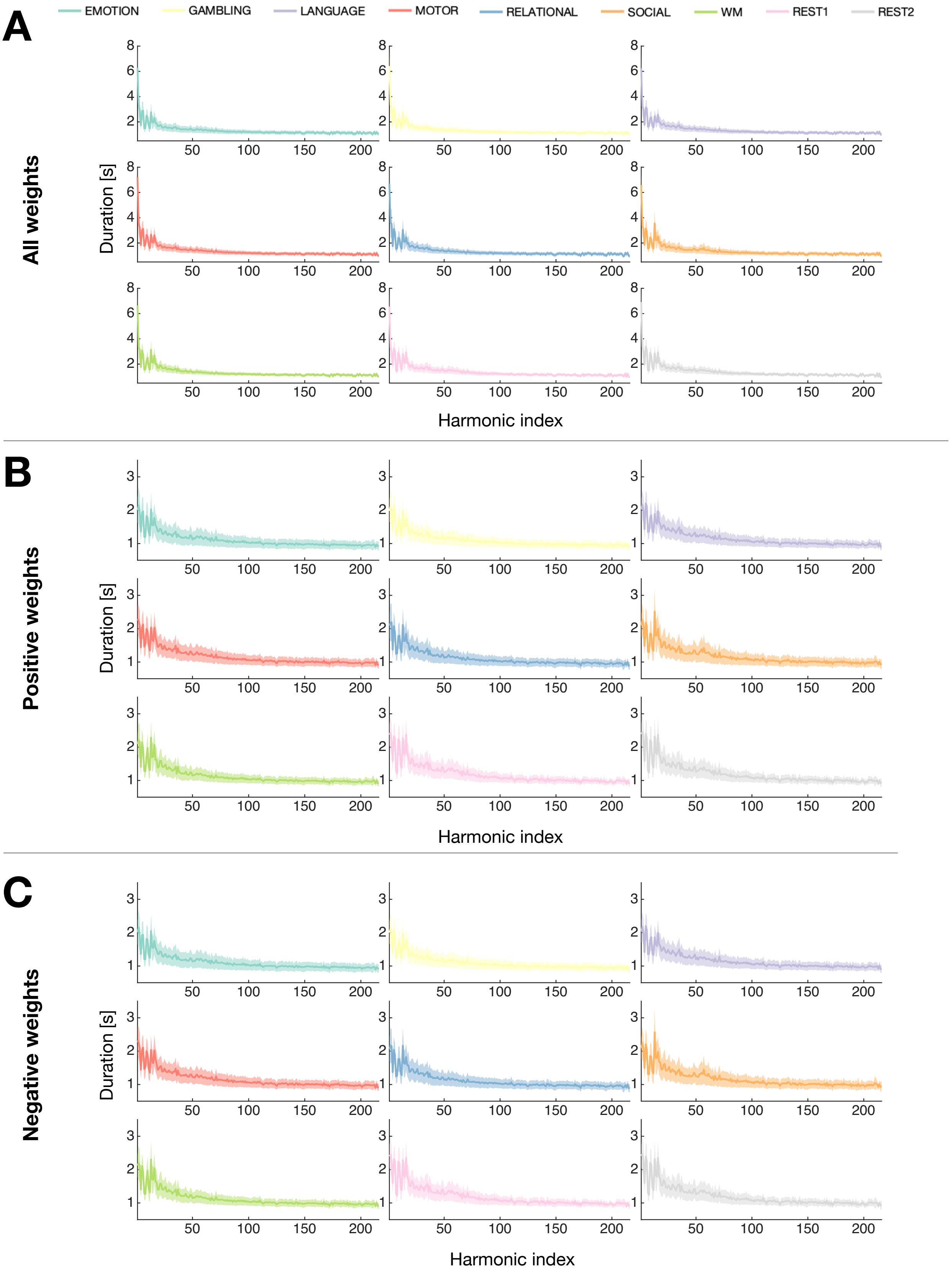

**Supplementary Figure 20: Spectral profiles of harmonic expression duration are preserved across paradigms and expression polarity.** Mean harmonic expression duration as a function of harmonic index for each HCP paradigm, computed using **(A)** all expression weights, **(B)** only positive weights, or **(C)** only negative weights. Shaded regions denote standard deviation across subjects. The spectral organization of harmonic expression duration is highly consistent across behavioral paradigms and remains highly similar when positive and negative expression weights are analyzed separately, indicating that the observed dynamics are not driven by a single expression polarity.

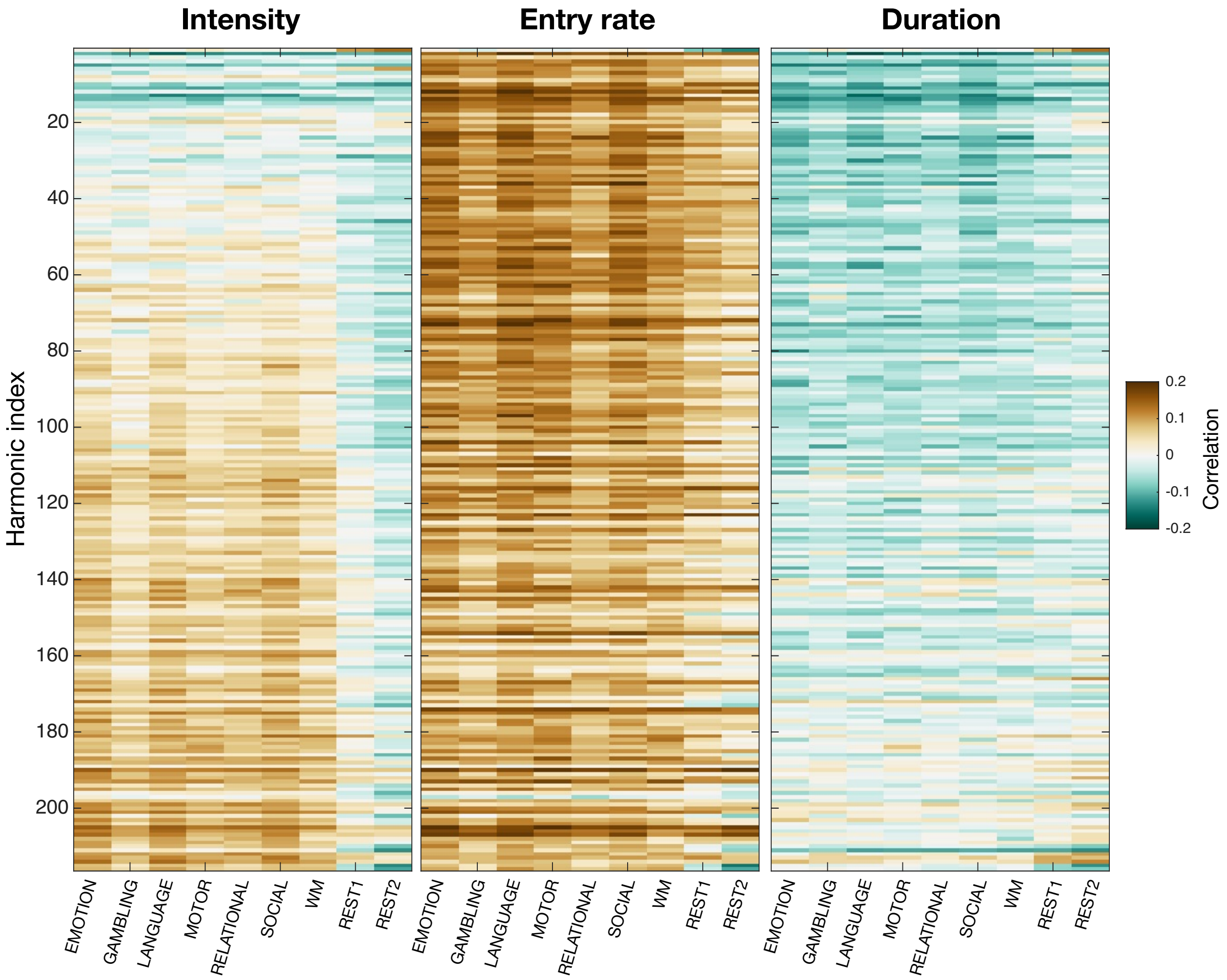

**Supplementary Figure 21: Harmonic expression dynamics exhibit only weak associations with head movement.** For each metric (each heatmap), harmonic (rows of a heatmap) and HCP paradigm (columns of a heatmap), Pearson's correlation coefficients reflective of the population-level association between mean framewise displacement and the metric at hand. Across harmonics and behavioral paradigms, head movement exhibits only weak associations with harmonic expression intensity, entry rate, and duration, indicating that the reported spectral organization is unlikely to be driven by motion-related artifacts.

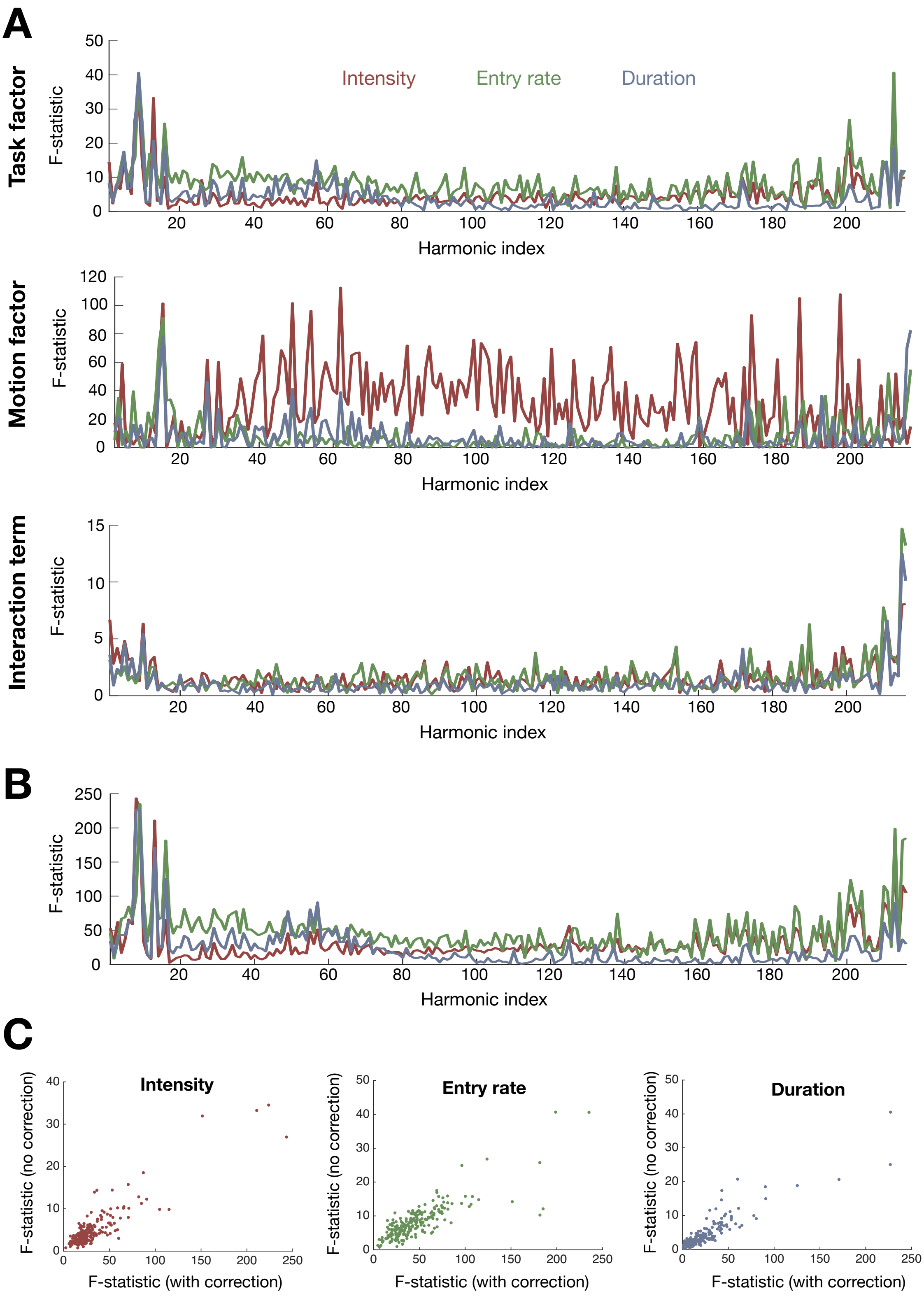

**Supplementary Figure 22: Head movement exerts only minimal impacts on cross-paradigm differences in harmonic expression dynamics.** For intensity (red), entry rate (green), and duration (blue), one-way ANOVA $F$-statistics for the task factor without accounting for head movement are shown in **(B)**, while two-way ANOVA $F$-statistics are shown in **(A)** for the task factor (top), the head movement factor (middle), and their interaction (bottom). **(C)** Comparison of the $F$-statistics obtained with (X-axis) and without (Y-axis) accounting for head movement for each metric. Together, these analyses show that accounting for head movement produces only minor changes in the task-related statistical profiles, with weak task $\times$ movement interactions restricted primarily to the lowest- and highest-frequency harmonics.

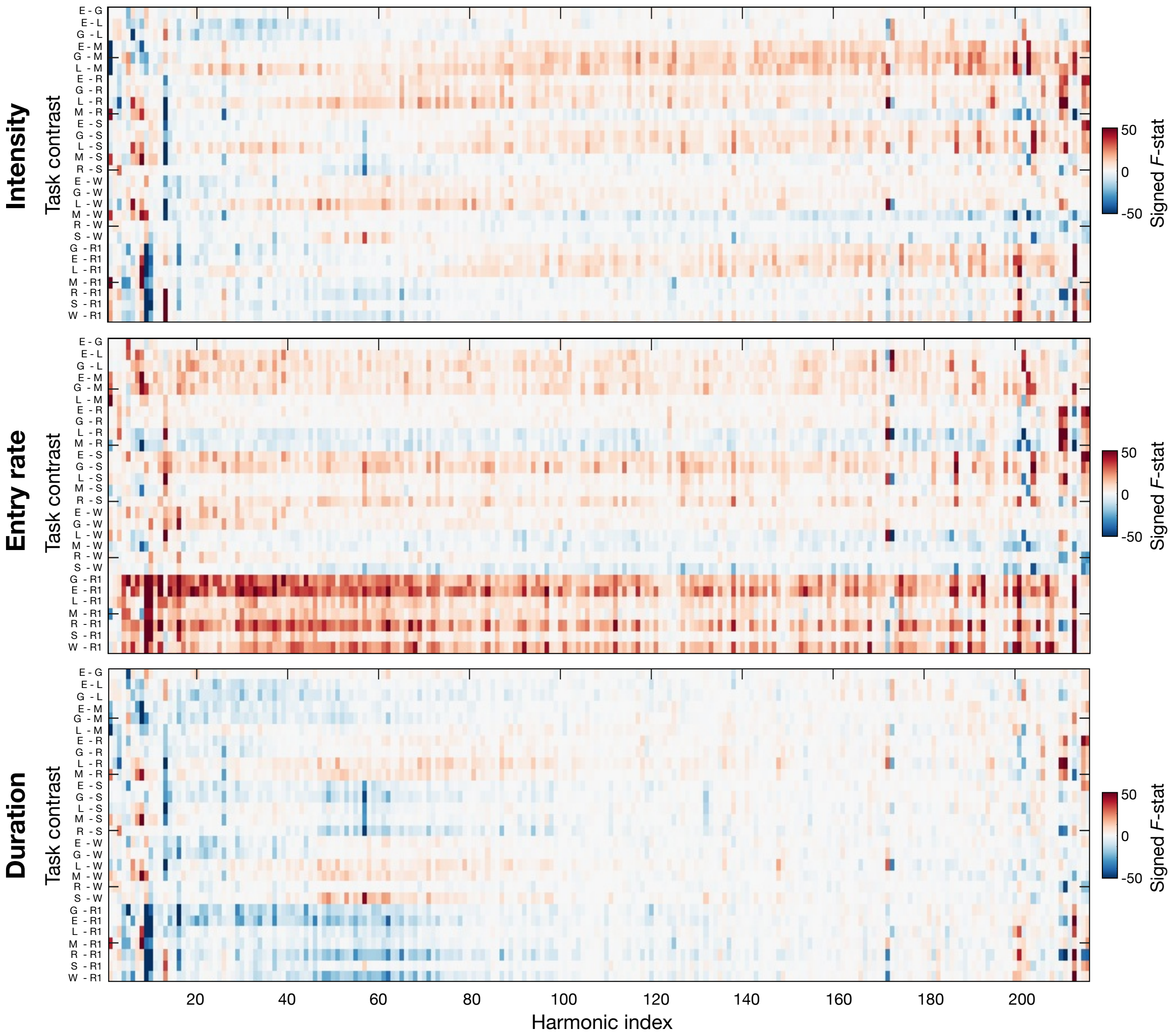

**Supplementary Figure 23: Metric-specific paradigm contrasts.** For intensity (top), entry rate (middle), and duration (bottom), signed $F$-statistics (the $F$-statistic multiplied by the sign of the corresponding contrast estimate) are shown for every task contrast (rows) and harmonic (columns). Positive values indicate larger metric values for the first paradigm of each contrast, whereas negative values indicate larger values for the second paradigm. These signed contrast maps constitute the complete set of pairwise comparisons that were summarized into the significance-based visualization shown in **Figure 4A**. E: EMOTION, G: GAMBLING, L: LANGUAGE, M: MOTOR, R: RELATIONAL, S: SOCIAL, W: WORKING MEMORY, R1: $\text{REST}_1$.

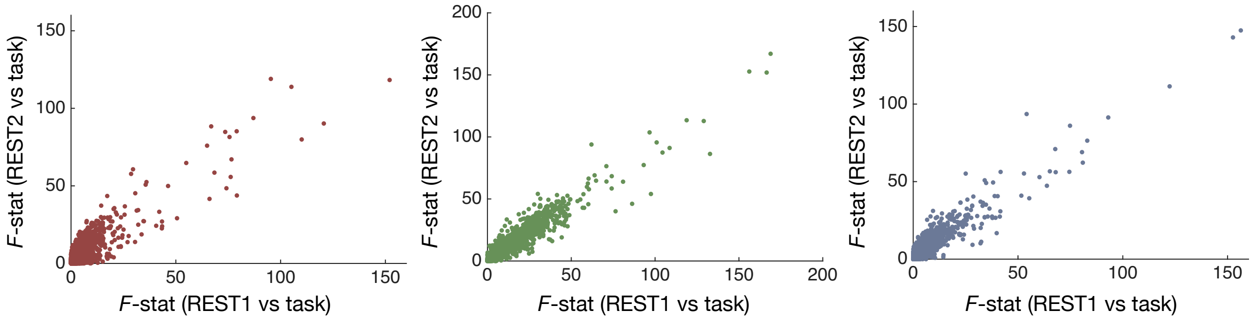

**Supplementary Figure 24: Rest-to-task contrast profiles are highly reproducible across independent resting-state acquisitions.** For intensity (left), entry rate (middle), and duration (right), F-statistics obtained for the $REST_1$-versus-task contrasts (X-axis) are plotted against those obtained for the corresponding $REST_2$-versus-task contrasts (Y-axis), using ANCOVA models that included mean framewise displacement as a continuous covariate. The strong positive associations demonstrate that rest-to-task differences are highly reproducible across independent resting-state acquisitions and therefore do not depend on the particular resting-state scan used for comparison.

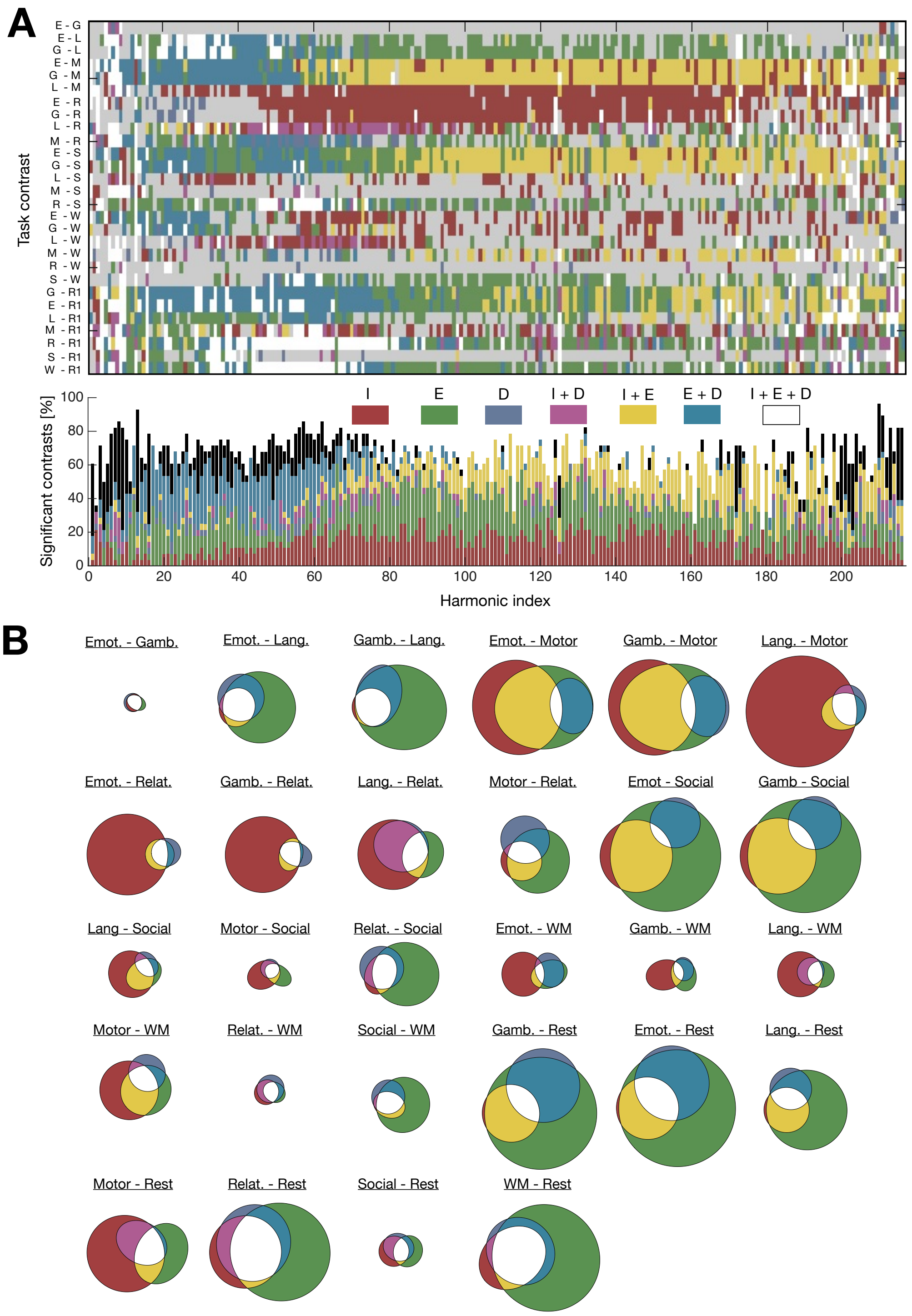

**Supplementary Figure 25: Patterns of cross-paradigm differences are preserved without accounting for head movement. (A)** (*Top*) For each task contrast (rows) and harmonic (columns), the color-coding indicates which combination of temporal metrics reached significance. Contrasts were evaluated using non-parametric rank-sum tests without including head movement as a covariate. Grey entries denote non-significant comparisons after Bonferroni correction. These results should be compared to those in **Figure 4A**, where head movement was explicitly modeled. (*Bottom*) Percentage of significant task contrasts for each harmonic, with color-coding the same as above. **(B)** Venn diagrams depicting the proportions of harmonics showing a specific combination of significant temporal metrics for each task contrast. The overall size of a diagrams is proportional to the percentage of significant harmonics. Together, these analyses show that the overall organization of significant cross-paradigm differences is largely preserved even when head movement is not modeled explicitly, indicating that the principal findings are robust to the statistical framework used.

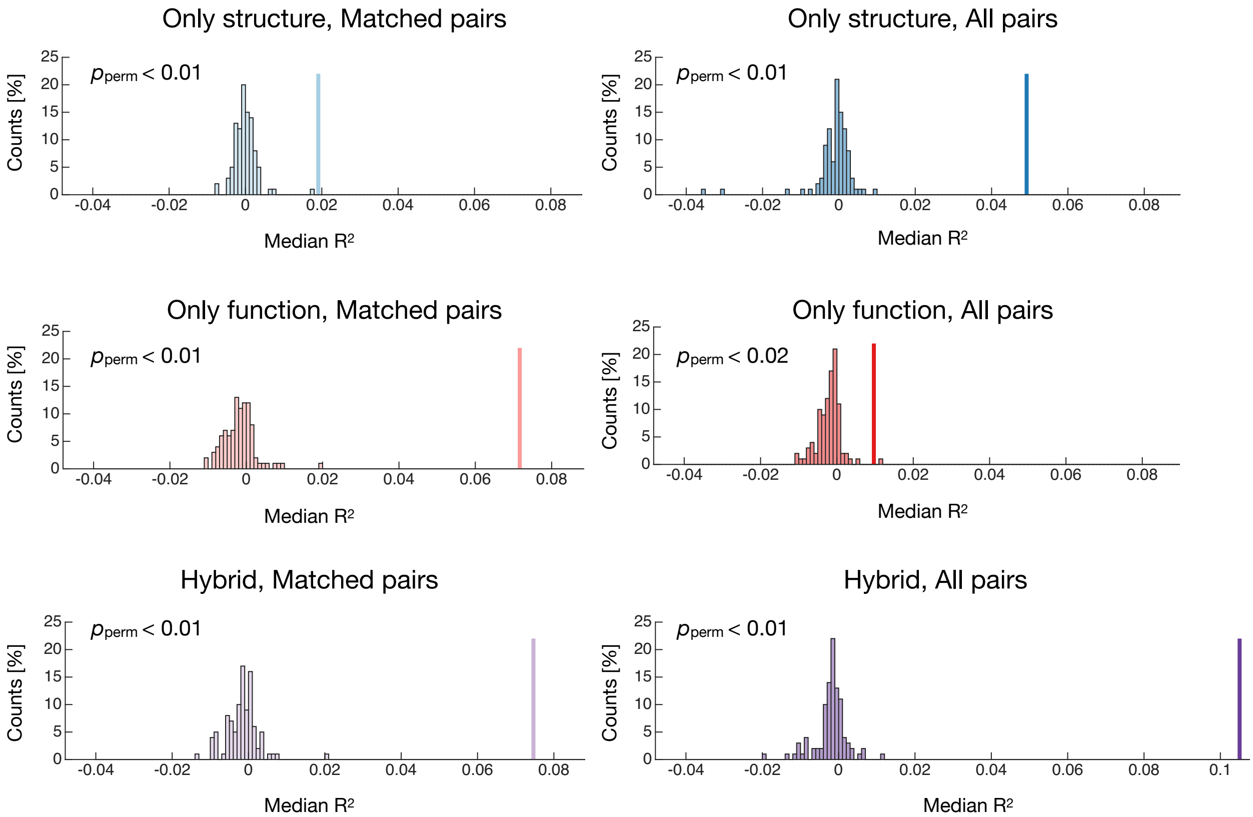

**Supplementary Figure 26: Permutation testing confirms above-chance prediction performance.** For all six models examined in **Figure 4B**, the observed median prediction $R^2$ obtained across nested cross-validation repetitions (vertical line) is compared with the null distribution obtained from permutation testing, after randomly permuting cognition factor scores across subjects and repeating the complete prediction pipeline (histogram). Color-coding is similar as in **Figure 4B**. In all six cases, the observed prediction performance significantly exceeded the permutation-derived null distribution ($p_{\text{perm}} \leq 0.02$), confirming that all prediction models performed above chance.

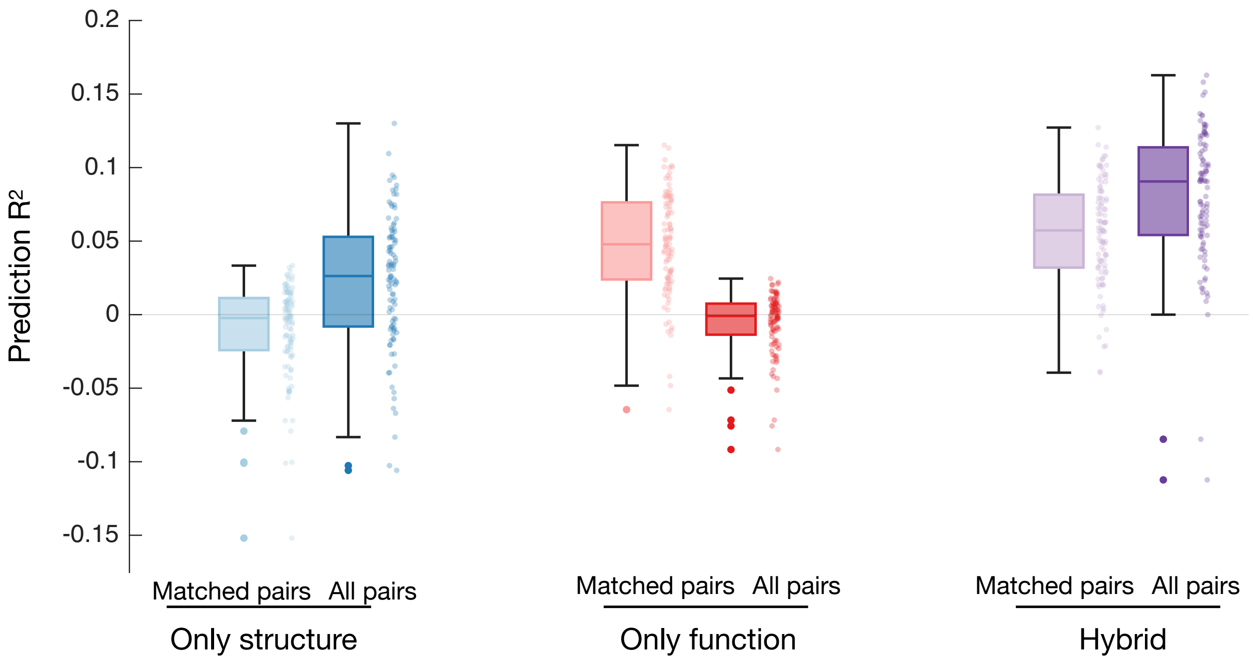

**Supplementary Figure 27: Prediction performance is insensitive to head movement.** Coefficient of determination $R^2$ for the prediction of *Cognition* using kernels constructed from structural similarity between harmonics (blue), functional similarity between cross-task harmonic power profiles (red), or their combination (purple), when head movement is not included as a covariate of no interest, in contrast to the analyses shown in **Figure 4B**. Light box plots correspond to kernels computed from matched harmonic pairs only, whereas dark box plots include all harmonic pairs. The qualitative similarity to **Figure 4B** indicates that accounting for head movement has only a minimal influence on prediction performance and does not alter the relative performance of the different kernel models.

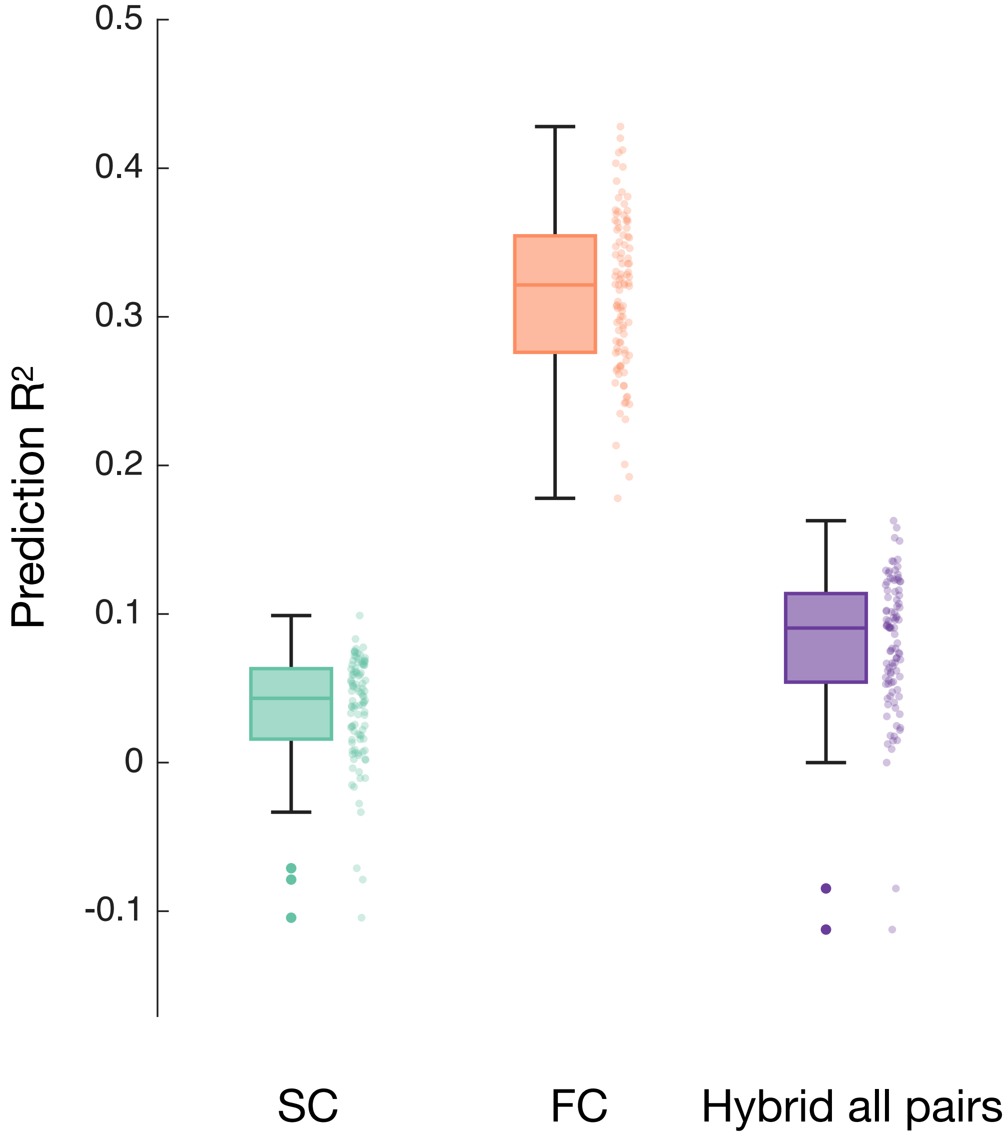

**Supplementary Figure 28: Prediction performance using standard structural or functional connectivity.** Coefficient of determination $R^2$ for the prediction of *Cognition* using kernels constructed from structural connectivity (turquoise), functional connectivity concatenated across paradigms (salmon), or the proposed hybrid kernel computed from all harmonic pairs (purple). Functional connectivity achieved the highest prediction performance, followed by the hybrid kernel and structural connectivity.
